# Multi-scale predictions of drug resistance epidemiology suggest a new design principle for rational drug design

**DOI:** 10.1101/557645

**Authors:** Scott M. Leighow, Chuan Liu, Haider Inam, Boyang Zhao, Justin R. Pritchard

## Abstract

A critical goal in drug discovery is the rational design of therapeutics that last longer in the face of biological evolution. However, this goal is thwarted by the astonishing diversity and stochasticity of mutational paths in the clinic. Beyond biophysical predictions of thermodynamics, we present stochastic, first principles models of evolution built on a large *in vitro* dataset that accurately predict the epidemiologic abundance of mutations to three different drugs across multiple leukemia clinical trials. Our ability to forecast the prevalence of resistance mutants required an understanding of the likelihood of the nucleotide substitutions that cause them. Beyond leukemia, a meta-analysis of acquired drug resistance across prostate, breast, and gastrointestinal stromal tumors (GIST) suggests that drug resistance in the adjuvant setting is significantly influenced by mutation bias. Our analysis points towards a new design principle in rational drug discovery: when evolution favors the most probable mutant, so should drug design.

## Introduction

Since *“On the Origin of the Species”* in 1859, the most powerful demonstration of natural selection is the pervasiveness of genetic resistance following the adoption of new drugs for viruses, prokaryotes, eukaryotes, and cancers^1–7^. Thus, efforts to rationally design new drugs that are less susceptible to evolutionary change are urgently needed.

Foundational stochastic models of evolutionary dynamics in cancer and infectious diseases have focused on the probability that most drug resistance mutations pre-occur in large populations of tumors, bacteria and viruses^1,8–12^. These theoretical arguments led to the practical insight that non-cross-resistant drug combinations are needed to combat genetic diversity. They also formed the basis for our current therapeutic regimens in HIV, TB, and cancer. However, despite this powerful example of evolution-guided therapeutic regimen design, drug resistance remains a problem. We posit that an important step forward involves using decades of improvements in evolutionary theory^1,12,21,13–20^ and population genetics^22–24^ to create additional design principles for drug discovery informed by evolution. We suggest that this might be achieved by expanding our ability to quantitatively predict which diverse resistance mutations can generate relapses in individual patients during treatment.

Recent and classic papers in cancer and bacteria have shown that biophysical methods and mutagenesis screens have great value in qualitatively identifying which mutations in a protein might lead to clinical resistance^25–27^. However, a long list of possible resistance mutations is challenging to incorporate into drug design. Which mutations will be most clinically abundant, and thus constitute “must hit” variants during drug development? To answer this question, we must go beyond qualitative predictions of possible resistant mutant identity to quantitative predictions of evolutionary outcomes. Recent progress in predictive evolution has shown that biophysical metrics can predict cellular fitness landscapes in antibiotic resistance^26^, and that fitness effects can forecast seasonal flu trends^28^. These exciting steps forward led us to ask an unanswered question in drug resistance research: Can the diversity and distribution of mutations that contribute to resistance evolution be quantitatively predicted? And could those predictions guide drug discovery?

There are two scales that combine to quantitatively determine which drug resistance variants arise across a population: 1) the host-level variables affecting *de novo* resistance generation and 2) the community level variables affecting the global spread of resistance. *De novo* variables include growth rates in the presence and absence of drug (which are often measured) as well as mutation rate, codon structure, genetic context and pharmacology (which are seldom measured). Human cancers offer a unique opportunity to investigate the process of parallel *de novo* resistance evolution in individual hosts because they lack community level variables that affect the spread of variants across a population (Figure 1A). Thus, predictive models of resistance to targeted cancer therapies are an interesting first test for predictive models of the evolution of drug resistance variants across real world populations.

**Figure 1:**
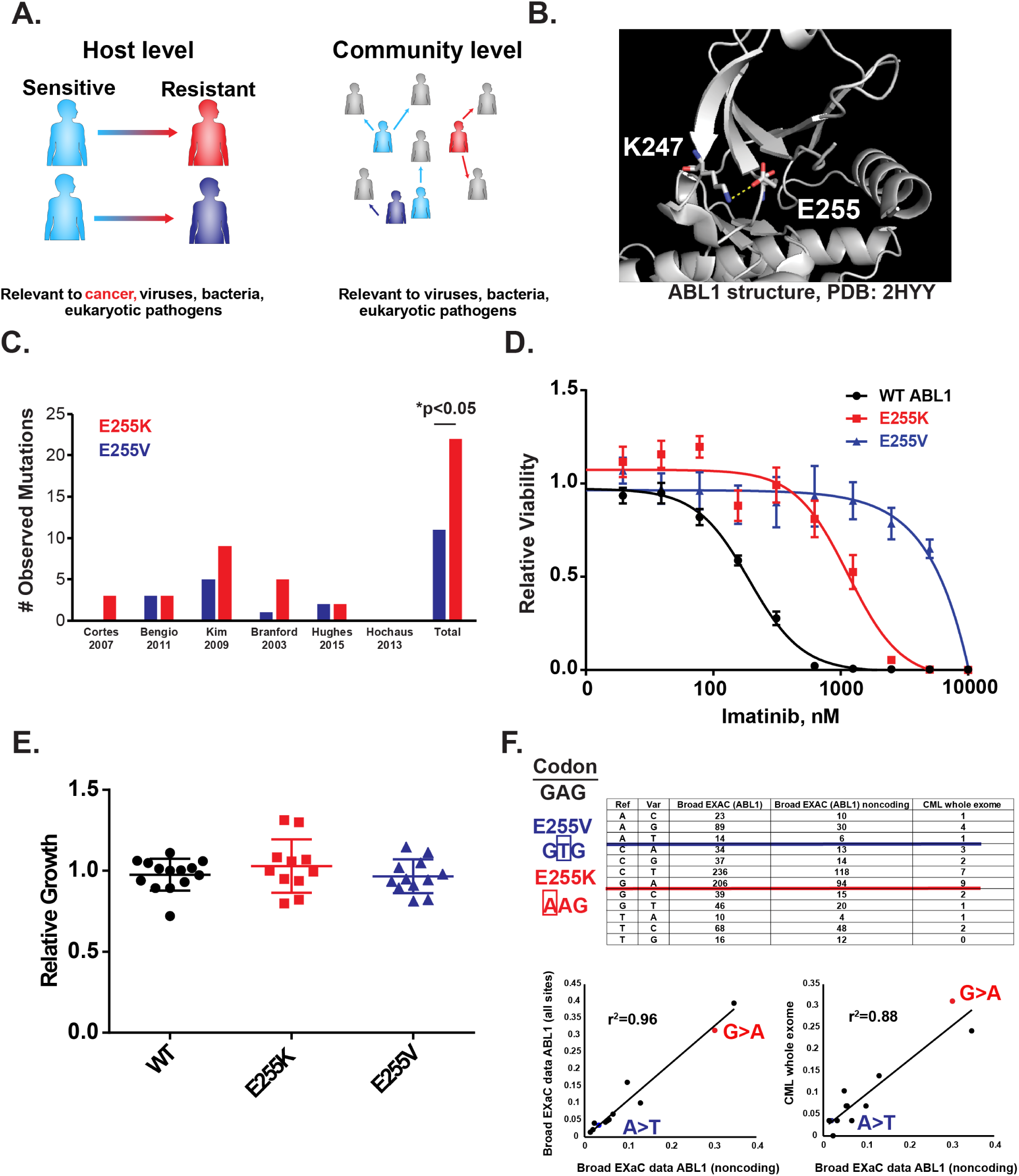
A salt bridge in ABL1 suggests that clinical abundance may not be predicted by the amount of drug resistance conferred. **(A)** A schematic of factors affecting the evolutionary dynamics of drug resistance. Without transmission, intra-tumoral variables are the only factors involved in cancer evolutionary dynamics and are limited to the host level. **(B)** ABL1 crystal structure. Ribbon diagram of secondary structure is shown. Image is zoomed in on the kinase P-loop. Loss of the E225-K247 salt bridge is associated with imatinib resistance. **(C)** Prevalence of E255K/V mutations in six imatinib clinical trials. A cross-trial sum is included, p-value is for chi-square test. **(D)** Imatinib IC_50_ curves for BCR-ABL transformed BaF3 cells. Relative viability is measured by Cell-Titer Glo relative to a DMSO control. N=3 per concentration, error bars are standard deviations. **(E)** Relative growth rates of BCR-ABL BaF3 variants. Each dot is an independent transduction and selection. N=11-14, error bars are standard deviations. **(F)** *Upper:* Codon structure of E255 and various measurements of nucleotide substitution frequencies. These measurements include counts of substitutions across the ABL1 gene identified in the Broad ExAC database; noncoding substitutions in ABL1 identified in the Broad ExAC database; and substitutions across the exome in CML patients. *Lower*: Different measurements of substitution likelihood were highly correlated.

The lack of a quantitative and predictive consideration of these evolutionary variables creates a gap in cancer patient care. In the targeted therapy of cancer, as first generation drugs are used and resistance liabilities are identified, drug discovery scientists have raced to develop second generation drugs that can target resistance mutations^29–32^. Second generation inhibitors can improve clinical outcomes in drug resistant and drug naïve patients, but vulnerabilities persist because patient need drives rapid drug development in the face of immature clinical data^29,31,32^. Thus, molecular design occurs before we know the true prevalence of specific mutations in the population. This means that solutions are offered in the clinic before the full scope of the resistance problem is understood. Structure-based drug design is now the industry standard to create potent second generation inhibitors and has succeeded in ABL1+ leukemias^30,33^, c-KIT/PDGFR mutated GIST^34^, ALK/EGFR+ lung cancers^35,36^, RET mutants/fusions^37^, and TRK fusions^38^. Rational design is typically based upon the biophysics of binding to the target and key counter targets, but not evolution. Thus, it can often identify whether a mutation can cause resistance, but it can’t determine how often we would expect to see that mutation in the clinic^39^. Using evolutionary theory to prospectively identify the residues and abundances that contribute to resistance following real-world drug use will improve pharmaceutical design. By developing a broader picture of drug resistance evolution before clinical data has matured, evolutionary criteria may be combined with structure-function analysis to guide next generation drug development.

In this study we parameterize stochastic, first-principles models of drug resistance. By systematically studying multiple variables that could affect *de novo* resistance generation, we show that predictive evolutionary modelling can forecast population patterns of drug resistance without requiring clinical measurements of resistant mutant specific resistance parameters. Moreover, we show that multiple treatment scenarios and biological architectures create populations that are sensitive to nucleotide substitution bias and codon path (together termed substitution likelihood) in the introduction of drug resistance variants in the clinic. We posit that next generation drug design could become more evolutionarily principled by adopting a simple design principle: when evolution favors the likeliest resistance mutation, so should drug discovery.

## Results

### A single salt bridge in ABL1 suggests that clinical abundance is not predicted by multiple fitness metrics

We first examined two drug resistance mutations identified in the literature that exist at the same amino acid in the BCR-ABL oncogene: E255. E255 forms a salt bridge interaction with K247, stabilizing the phosphate binding loop (P-loop) that clamps over the ATP competitive inhibitor in the active site of the molecule (Figure 1B). E255 can mutate to become either E255V or K, both of which abolish the charge interaction and promote clinical resistance to imatinib (Figure 1C), a BCR-ABL inhibitor used to treat patients with chronic myelogenous leukemia (CML). Interestingly, tallying the abundance of E255V and E255K mutations across clinical studies suggested that E255K was more prevalent (21 patients) than E255V (10 patients) (p<0.05, chi-square test).

Next, we sought to examine why E255K might be more prevalent than E255V. We made BaF3 cells that harbored wild type, E255V, and E255K BCR-ABL (a common model for BCR-ABL targeted therapies^40^), and we examined their relative sensitivity to imatinib. As expected, both mutations provided resistance to imatinib, but E255K provided significantly less resistance than E255V (Figure 1D). Importantly, this difference in phenotype occurs at clinically relevant concentrations of imatinib, which has a C_max_ of 5.3 μM and a C_trough_ of 2.4 μM at steady state^41^. While further investigation into the biophysics of P-loop dynamics could reveal the molecular mechanism behind this difference in drug resistance, we aimed to understand how a mutation that is worse at conferring drug resistance (regardless of the biophysical mechanisms) might become more prevalent in humans. Because the less drug resistant variant appeared to grow out more often, we examined the relative growth rate of E255V and K in the absence of drug. If E255V grows significantly slower before imatinib exposure, that might help to explain its lower incidence in a human population. However, the relative growth rates for E255V, E255K and wild type BCR-ABL cells were indistinguishable (Figure 1E), despite having 80% power to detect a ∼ 10% growth rate difference (see GitHub).

Thus, we hypothesized that a mutation from E>K might be more likely to occur than a mutation from E>V. Examining the genetic code, E>K requires a transition from G>A, while mutating from E>V requires a transversion from A>T. No other single nucleotide substitutions can cause these two mutations. While transitions are usually more likely than transversions, mutation biases are known to vary across genes and cancer types. We sought a direct measurement of the mutation bias in ABL1 and CML. To do this, we turned to multiple data sets that have analyzed mutations in cancer and variation in the normal human exome. As an example, the Broad ExAC data set of 100,000 human exomes (see Methods, Table S1) can be used to estimate ABL1 mutation bias by focusing on the extremely rare variants that constitute 90% of the variation in the ExAC data. Using this data set we examined rare (<1/10,000) synonymous mutations, missense mutations, and noncoding mutations in the ABL1 gene (Figure 1F, upper). Splitting these mutations by nucleotide type on the transcribed strand to account for biases due to transcription, we quantified the nucleotide substitution biases in the ABL1 gene. The measured bias was consistent with a transition-transversion bias, i.e. G>A mutations were much more likely than A>T mutations. Moreover, it was largely invariant depending upon the exact mutation types utilized (synonymous, nonsynonymous, intronic) and consistent with previous literature. Finally, this ABL1 mutation bias also highly correlated with the mutation biases that were measured across the CML exome (Figure 1F, lower).

We observed that the substitution likelihood of a given amino acid mutation (defined as the nucleotide substitution bias and the individual codon path), but not the fitness in the presence or absence of drug, is correlated with the clinical prevalence of E255V versus E255K.

### An analytical model of stochastic dynamics identifies parameter regimes in which the likeliest mutations can predominate

Because E255K was more likely to occur than E255V, but not more fit in the presence or absence of drug treatment, we asked whether it makes theoretical sense that a more likely mutation can beat a more resistant one to create *de novo* drug resistance.

To study this, we developed a probability model for the stochastic evolutionary dynamics of a hypothetical drug target with two possible resistance alleles. Allele A is more resistant (more fit in the presence of drug) but less likely; Allele B is less resistant but more likely (Figure 2A). The probability that either allele drives relapse was calculated from the allele-specific probabilities of seeding a detectable resistance clone. In cases where both resistance clones of the two alleles are present, the first allele to reach clinical detection was considered to dominate relapse. This “race” to detection was formulated as two steps: mutation and growth. While Allele A is expected to grow out faster, Allele B is expected to mutate first, giving it a head start (Figure 2B). From this model, we derived an asymptotic solution for the probability that either allele drives relapse (Appendix S1).

**Figure 2:**
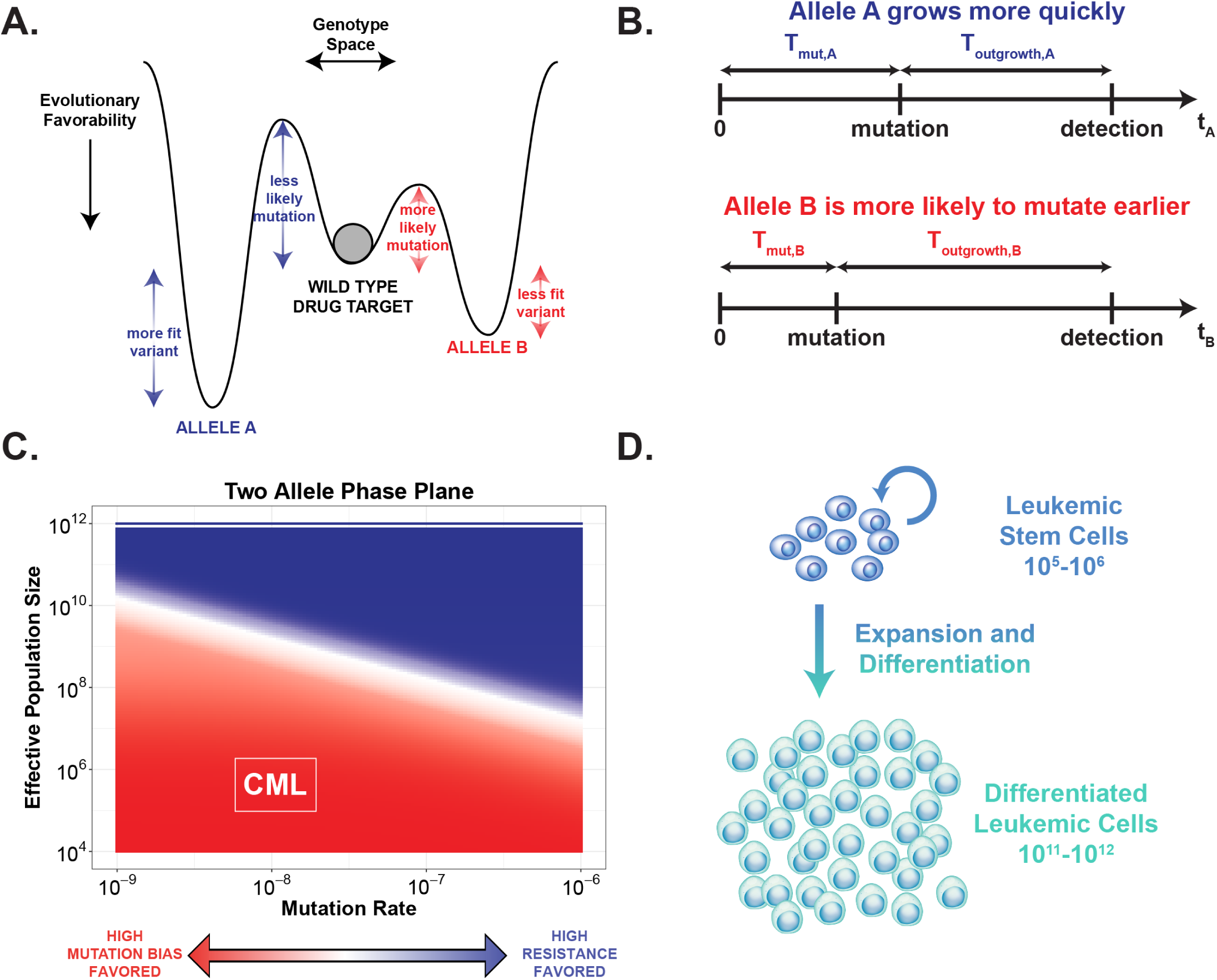
An analytical model of stochastic dynamics identifies where survival of the likeliest can occur. **(A)** Evolutionary landscape for a theoretical drug target gene with two potential resistance alleles. Allele A is assigned a high fitness and low probability; Allele B is assigned a low fitness and high probability. **(B)** A schematic of a general timeline for mutation and outgrowth for either allele given the assigned evolutionary profiles. In cases where both mutations occur, the first resistant clone to reach detection drives relapse. **(C)** A phase plane of the results of our probability model across many mutation rates and effective population sizes. Color indicates whether Allele A (**dark blue**) or Allele B (**red**) is more likely to drive relapse. In regions where Allele B is more dominant, we expect mutation bias to be a primary evolutionary force. **(D)** Schematic of general leukemic cell population hierarchy. Only mutations in leukemic stem cells can form stable resistance clones, effectively limiting the population size to 10^5^-10^6^.

We then evaluated these probabilities across a region of the effective population size/mutation rate parameter space to identify parameter regimes where Allele B (the less fit, more likely allele) is more likely to drive relapse. We refer to this phenomenon as survival of the likeliest, and it is strongest in the region of the phase plane with low mutation rates and a small effective population (Figure 2C), corresponding to populations with low heterogeneity. Here, where the opportunity for mutation is limited, unlikely mutants may not spawn at all, regardless of their fitness. Thus, under these conditions, it is more important for a resistance mutant to be likely than to be fit. In contrast, as mutation rate and effective population size increase, Allele A is more likely to dominate, reflecting a more standard survival of the fittest model. Thus, the degree of heterogeneity, as shaped by population size and mutation rate, determines when substitution likelihood can play a driving role in drug resistance epidemiology.

These theoretical results point towards the correlation of mutation and codon biases observed in real-world CML resistance incidence. The population structure of CML includes a well-characterized leukemic stem cell population of 10^5^-10^6^ cells that gives rise to peripheral differentiated leukemic cells (Figure 2D). This hierarchy effectively restricts the population size, since only resistance mutations that occur in leukemic stem cells have any clinical consequence.

Thus, CML can be placed in the “low heterogeneity” regime of the parameter space. Together, theory and empirical data support the idea that low heterogeneity populations are most strongly influenced by mutational likelihood.

### Epidemiologic incidences of ABL1 resistance mutations are best predicted by how likely they are

To approach this problem more comprehensively, we gathered data from hundreds of parallel clinical evolution experiments in BCR-ABL+ leukemias across four continents over 17 years (Data S1)^42–47^. Specifically, we identified clinical trials with clear clinical resistance criteria and codon level resolution. 268 high confidence clinical cases of imatinib resistance were identified in these studies. Tallying mutations at individual amino acid positions, we found that the 19 most abundant amino acid mutations account for ∼ 95% of the resistance mutations identified (Figure 3A). Notably, 85% of the resistance prevalence captured by this set of mutations is caused by a mutation other than the fittest (T315I; imatinib IC_50_ = 8711 nM).

**Figure 3:**
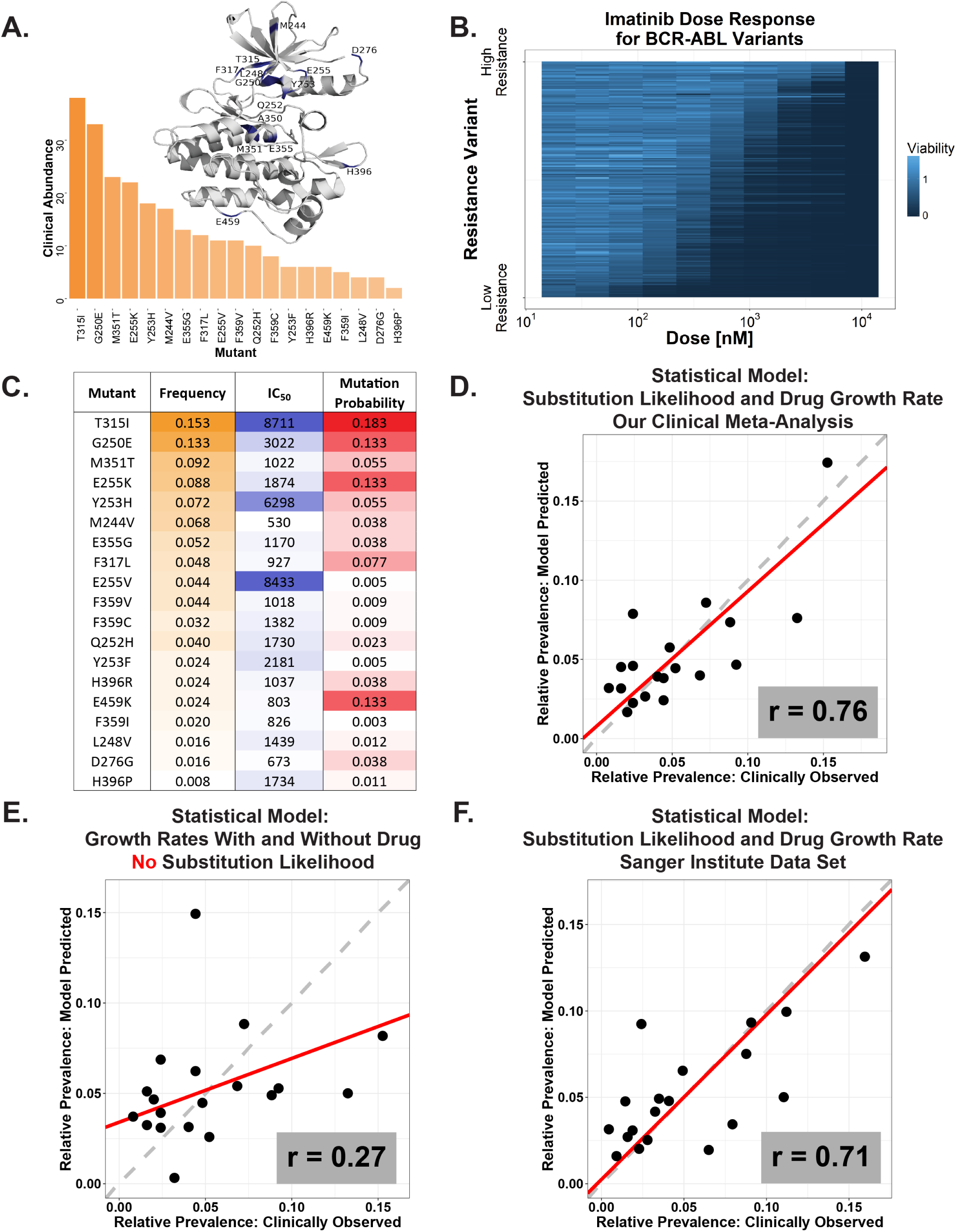
Epidemiologic incidences of ABL1 resistance mutations are best predicted by how likely they are. **(A)** A crystal structure ribbon diagram of the ABL1 kinase domain and distribution of the 19 most prevalent BCR-ABL resistance mutants. These 19 variants account for approximately 95% of resistance mutations observed clinically in the six studies from Figure 1C. **(B)** Drug-dose response was measured by Cell-Titer Glo and is plotted in the heatmap. N=3 independent infections for all 20 cell lines (WT and mutants). Each BCR-ABL BaF3 line was dosed with 11 serial dilutions of imatinib in triplicate. All cell lines are ordered by sensitivity. Raw data and code are available on GitHub. **(C)** A table that includes values used to build regression models: the frequency of each resistant mutant as determined by our clinical meta-analysis, imatinib IC_50_s (in nM) normalized for genetic background, and substitution likelihood calculated from analysis of the Broad ExAC data in Figure 1F. **(D)** Observed versus predicted plot for a regression model of clinical mutation prevalence (determined by our clinical meta-analysis) regressed against growth rate in the presence of drug and substitution likelihood (which is the final model). Points are specific ABL1 mutations; x-values are their observed frequency and y-values are their frequency predicted by the model. Pearson correlation for observed and predicted prevalences is r = 0.76. **(E)** Observed versus predicted plot (as in Figure 3D) for the regression model of clinical mutation prevalence built only on the growth rates in the presence and absence of imatinib. Pearson correlation r = 0.27. **(F)** Observed versus predicted plot (as in Figure 3D) for independent Sanger Institute data set. Pearson correlation r = 0.71.

For each of these 19 amino acids, we generated three independent isogenic BaF3 cell lines and measured the IC_50_ of imatinib in each cell line in triplicate across 11 serial drug dilutions. (Figure 3B). While this set of 19 clinical point mutations in ABL1 is the most systematically assembled *in vitro* data set to date (regarding clinical abundance), many of these mutations have had their IC_50_s measured in other resistance studies in other labs. Thus, the literature presents us with an opportunity to estimate how genetic context and experimental conditions might affect drug response. Supplemental Figure S1 shows that cross study correlations are high (Pearson’s r∼ 0.9 or higher for four studies), but systematic shifts in slope exist between studies. This might suggest that the genetic drift in cell lines in an individual lab can influence drug potency and is consistent with recent findings^48^. Thus, we have an opportunity to account for the effects of genetic background upon drug resistance. To do this, we normalized all systematic differences between the data sets into one “mean” data set and combined all the data from the literature with our data (see Methods and Data S1). This cross-study approach leverages the genetic drift across cell lines to get the best estimate of the average isogenic effect of BCR-ABL mutations.

Beyond genetic background normalized IC_50_ measurements, we also measured the drug-free growth rates of all ABL1 mutants from 6-10 independent lentiviral infections (Figure S2A). Many independent infections were critical because the growth rates of individual clonal selections exhibit high variance. We also measured the substitution likelihood of a given amino acid conditioned upon a resistance mutation event (Figure 3C, see Methods and GitHub). Negative binomial regression was used to predict the incidence of individual resistance mutants across the human population (Supplemental Figure S2B, R-markdown file on GitHub). After considering all possible single variable models, we built multivariate models, identifying the best N-variable model by leave-one-out-cross validation (LOOCV) (Figure S2C). A two-variable model built on normalized IC_50_ values and substitution likelihood (Figure 3D) fit the data well, exhibiting a Pearson correlation between model-predicted and observed values of r = 0.76. This model outperformed one built on normalized IC_50_ values and drug-free growth rates (Figure 3E), with a correlation of r = 0.27. While the statistical model suggested that the amount of drug resistance and the substitution bias were both significant and predictive (Figure 3D/E and Figure S2C, R-markdown file in GitHub), substitution likelihood was the more significant predictor of clinical abundance (p=3.3e-5 for substitution bias vs p=0.016 for IC_50_). To verify that our result wasn’t overfit, we also identified a likely independent data set of the abundance of different ABL1 mutations that is curated by the Sanger Institute. While these mutations have a less clear clinical provenance than our analysis, we decided to use them as a “test set” because they had considerably more data. The same two variables performed well in this independently generated model (Figure 3F, Supplemental Figure S2D). However, in case our data has any overlap with the Sanger Institute data, we also verified that the results for substitution likelihood were similar when our counts were subtracted from the Sanger Institute counts (p<0.0001 in the same bivariate regression model). The reproducibility in a second data set highlights the significance of substitution likelihood as the most important predictive variable.

Thus, substitution likelihood, rather than drug resistance, is the more significant variable predicting the relative abundance of the 19 mutants that account for 95% of the clinical mutations in BCR-ABL for patients with CML.

### A stochastic, first principles, multi-mutation model of three inhibitors predicts the clinical prevalence of resistance mutations across ABL1

Our experiments and epidemiologic analysis indicated that an understanding of substitution likelihood is necessary to predict the clinically-observed distribution of specific resistance mutations. Because of this, we wanted to know if we could predict the frequency of mutations from a mechanistic model built from first principles.

To do this, we first identified a simple, clinically-parameterized model of CML treatment that incorporates hematopoietic stem cell division and differentiation^49^. To adapt the system to our question, we added our 19 resistance variants of interest to the model (Figure 4A). Each mutant was parameterized with an allele-specific substitution probability and a drug kill term. The drug kill rates measured in our *in vitro* experiments were linearly scaled by the ratio of net growth rates of BaF3s and *in vivo* LSCs, under the assumption that the relative size of the resistance phenotype was preserved across the *in vitro* and *in vivo* systems. We used cell-based measurements in the presence of human serum to appropriately scale drug exposures to the effective *in vivo* levels (see Methods and Appendix S2).

**Figure 4:**
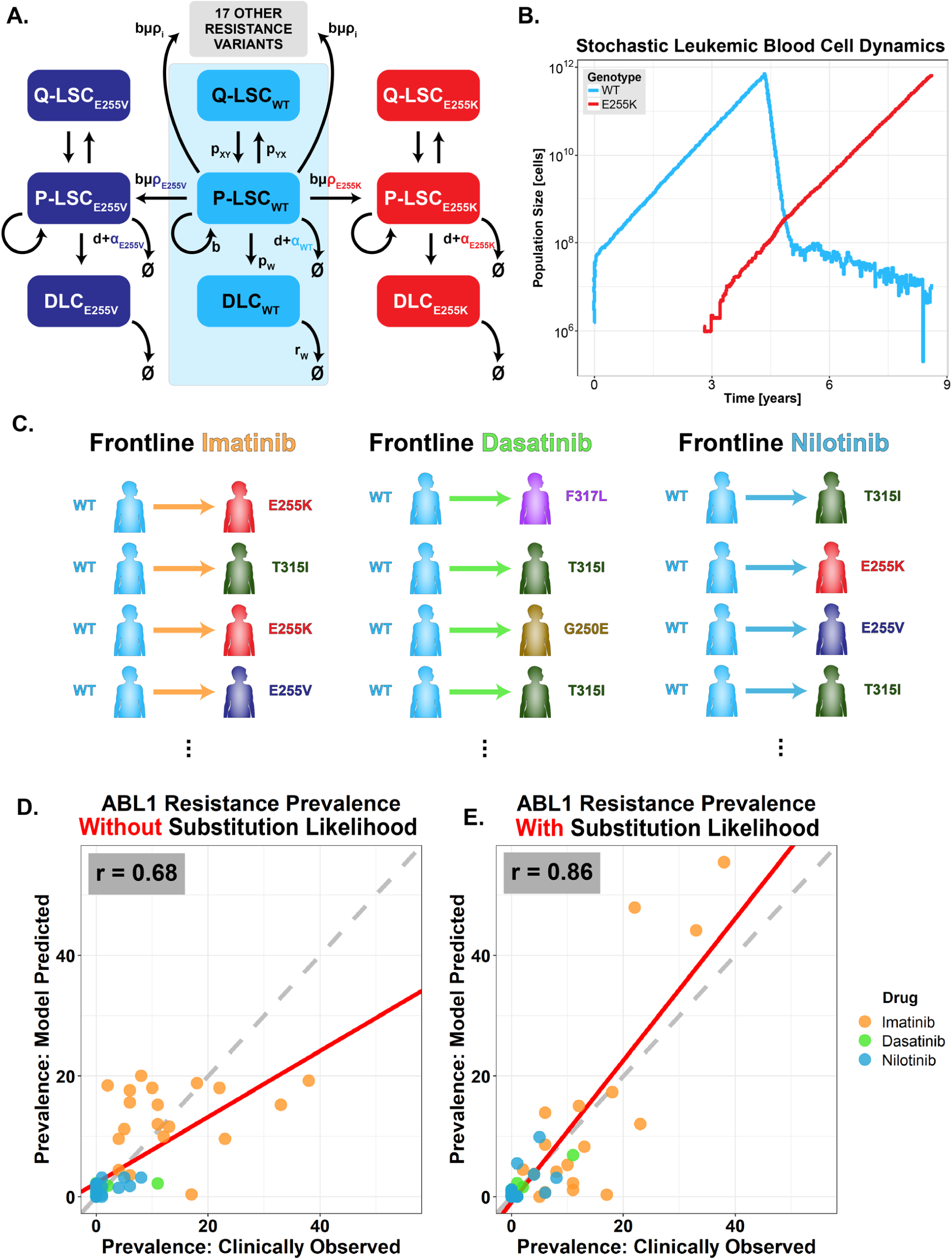
A stochastic, first principles, multi-mutation model of imatinib treatment predicts the clinical prevalence of resistance mutations across ABL1. **(A)** Schematic of stochastic CML evolutionary dynamic model. The initial deterministic model of three differential equations (shaded in **blue**) is from Fassoni et al. 2018^38^ and is fit to phase 3 clinical data. We reformulated a stochastic version of 60 differential equations parameterized from Fassoni et al. and our clinical data. Leukemic stem cells alternate between proliferating (P-LSC) and quiescent state (Q-LSC). P-LSC give rise to differentiated leukemic cells (DLC). P-LSCs may also spawn a resistant subclone P-LSC_i_ when dividing. The allele-specific mutational probability is given by ρ_i_. Note that we added the ability for all 19 resistance mutations to occur, such that there are 20 sets of differential equations with three populations per mutant. The system is solved stochastically. **(B)** An example stochastic simulation of the model described in Figure 4A. **(C)** Simulations were conducted (as in Figure 4B) 10,000 times for each of three BCR-ABL inhibitors: imatinib, dasatinib, and nilotinib. Resistance alleles were tallied across simulations for the three drugs. **(D)** Simulation results for the stochastic model without mutation bias (uniform ρ_i_). The Pearson correlation between observed and predicted prevalences is r = 0.68. **(E)** Simulation results for the stochastic model with mutation bias (allele-specific ρ_i_). The Pearson correlation is r = 0.86.

Importantly, this methodology does not rely on fitting the model to the prevalence of specific resistance mutations. Therefore, it only requires a mechanistic model of treatment (i.e. birth/death rates), a list of candidate mutations (which could be generated from structure-based design or mutagenesis), measurements of substitution likelihood and a tractable *in vitro* system to measure the effects of putative resistance mutations.

We simulated this system of 60 differential equations stochastically (Figure 4B) for 10,000 virtual CML patients treated with imatinib (see Methods). Stochasticity allows the potential for random mutation to seed each resistant variant. Each patient was assigned a pharmacokinetic profile from clinically-observed distributions of *in vivo* drug concentration^50^. Similarly, patient-specific tumor detection sizes were drawn from real-world distributions^51^ (Appendix S2). Across these 10,000 simulations, we totaled the number of *in silico* patients that relapsed with each mutation.

We repeated this process of measuring IC_50_s for all 19 resistance variants, simulating pharmacokinetic profiles, and parameterizing an evolutionary model, for second-generation BCR-ABL inhibitors nilotinib and dasatinib. Both TKIs have been previously evaluated in frontline clinical trials. This allowed us to appraise our predictive model with *in vitro* data for a total of three clinically-evaluated drugs (Figure 4C).

Importantly, we ran simulations for a null model without substitution likelihood, where individual amino acid changes were assigned the same probability, and compared its predictive value to that of a model parameterized with substitution likelihood. In examining the distribution of mutations across the ABL1 kinase for imatinib, nilotinib, and dasatinib, a model that considered substitution biases was required to accurately predict the abundances of individual mutations (Figure 4D/E; Pearson’s correlation r=0.68 without substitution likelihood vs r=0.86 with).

While our model was quantitatively accurate, clinical data for resistance to frontline nilotinib and dasatinib suffers from low counts (0-3 counts in most cases) and wide confidence intervals. Thus, to more completely evaluate our models’ predictive power in the context of these two drugs, we also examined categorical accuracy, i.e. the ability to say whether a mutation should or shouldn’t show up in a clinical trial. To do this, we downsampled our dasatinib and nilotinib simulations to reflect the size of the frontline trial. These virtual clinical trial results informed a binary classifier model – if a mutation was present, it was categorized as resistant; if not, sensitive. We repeated this for 10^3^ simulations, each time constructing an ROC curve to quantify the accuracy of the binary classifier against the real-world clinical data. Averaging the ROC curves across all simulations (Supplemental Figure S3A/B), we found that substitution likelihood considerably improved the models’ ability to classify a variant as likely to occur as a resistance mutation (AUC = 0.79 without mutation bias vs 0.91 with, for dasatinib; AUC = 0.65 vs 0.82, for nilotinib). Given that these predictive models were not fit to any patient data, they could be used to forecast which resistance mutations are identified in any CML clinical trial *a priori*.

These results demonstrate that a first principles, mechanistic model can predict the distribution of mutations seen in the clinic if it contains parameters that account for substitution likelihood. This improvement is consistent across imatinib, dasatinib, and nilotinib and underscores the importance of substitution likelihood in these models. To our knowledge, no previous mechanistic model has ever quantitatively predicted the epidemiologic diversity of clinical resistance mutations.

### Evolution-guided drug design could inform principled decisions between mutational vulnerabilities during drug development

To further investigate how an evolutionarily-informed approach to drug design might affect the clinical prevalence of resistance, we conceived of a hypothetical BCR-ABL TKI, which we call “maxitinib”, designed with mutational liabilities in mind. Given the same number of mutational vulnerabilities in a target (here we use five), maxitinib would be designed via structure-based drug design to target the five most likely mutants. If this could be achieved, how would that alter the overall incidence of resistance that arises in the clinic?

To test this, we simulated a cohort of *in silico* frontline CML patients for several target profiles of our hypothetical drug maxitinib; we name these distinct versions of maxitinib K1 to K15. Each of the hypothetical drugs K1-K15 denote a version of maxitinib that was designed to target a different set of five of the 19 previously-discussed imatinib-resistant mutants. The top five most likely mutants are sensitive to maxitinib K1; the second through sixth most likely mutants are sensitive to maxitinib K2; and so on (Figure 5A, see Methods and GitHub for model description).

**Figure 5:**
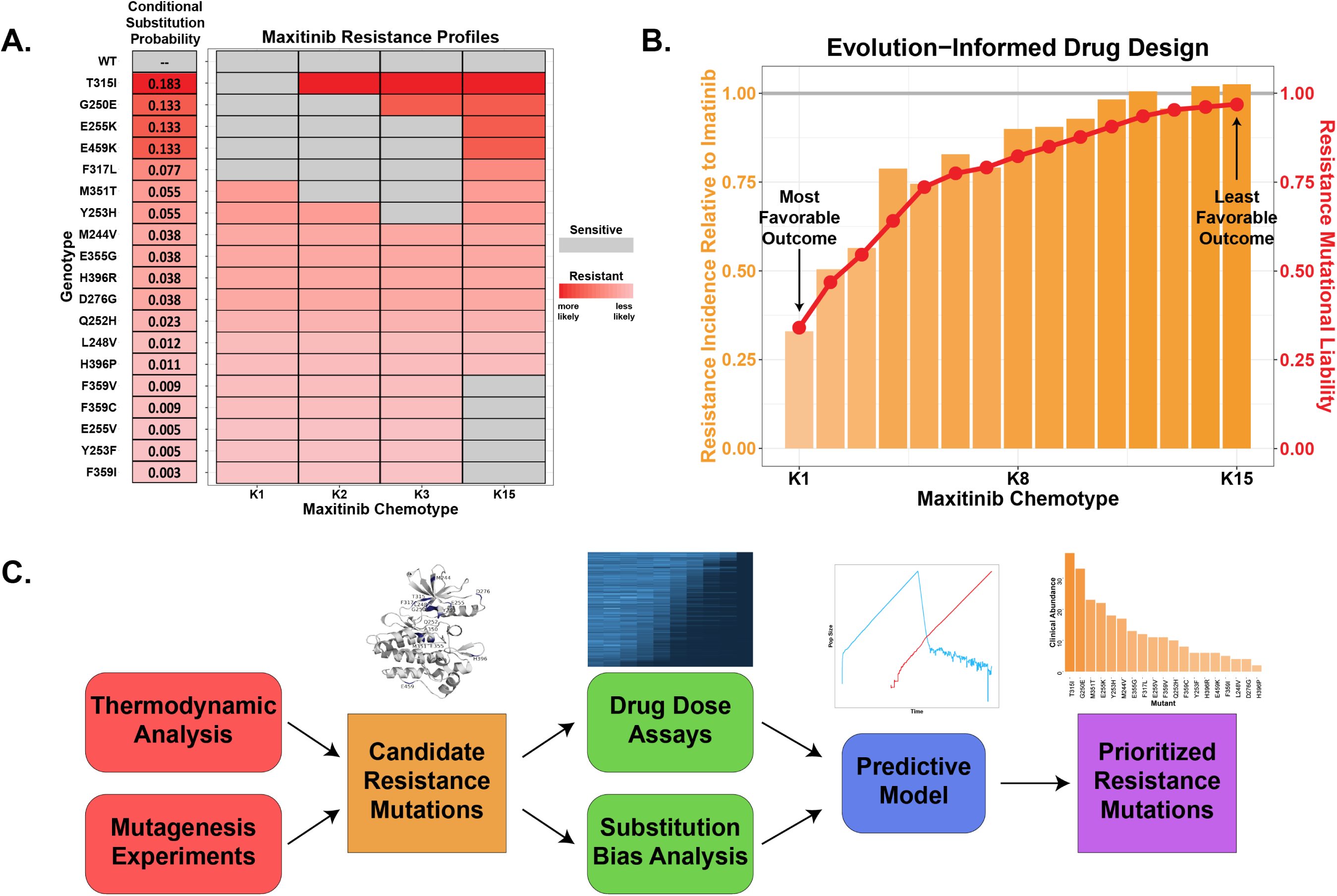
An evolution-guided approach drug design is predicted to minimize resistance prevalence. **(A)** Resistance profiles for versions of a hypothetical drug, “maxitinib”. Maxitinib K1 targets the first through fifth most likely mutants; K2 targets the second through sixth most likely; and so on. **(B)** Maxitinib simulation results. **Orange** bars represent the resistance incidence of each maxitinib chemotype relative to imatinib. **Red** points indicate mutational liability, defined as the sum of conditional substitution likelihoods of mutations that confer resistance to each chemotype. **(C)** A proposed workflow for evolution guided drug design. Potential resistance mutations could be generated by structure driven simulation, unbiased mutagenesis, or CRISPR base editors. Mutations can then be analyzed as single IC_50_s or in a pooled format. Indication specific information on the substitution biases would require the mutational signature for a given disease. Fitness and bias estimates could then be coupled into a mechanistic model of drug resistance that would predict the clinically most abundant resistance mutations.

The model predicts a 67% reduction in resistance incidence relative to imatinib for maxitinib K1, the hypothetical chemotype that targets the five most likely mutants (Figure 5B). This reduction is larger than either nilotinib or dasatinib (even though there are at least as many mutations that confer resistance to maxitinib K1 as nilotinib or dasatinib), and predicts that the evolutionarily optimal chemotype would prevail in the clinic.

Considerations relating to structure, conformation, and intermolecular interactions complicate drug design and would sometimes preclude the development of a drug that targets exactly the N most likely resistance mutants. However, as small molecules are screened, and tradeoffs are identified, evolutionary models could choose a molecular profile that predicts less resistance when used in the clinic. Our models suggest that designing molecules in this way could create second generation inhibitors that minimize drug resistance (Figure 5C).

### Evolution of resistance to drugs used in the adjuvant setting is shaped by substitution likelihood

Figure 2 identifies conditions where predictive evolutionary modeling would benefit from measurements of substitution likelihood. While CML is one example where the survival of the likeliest plays a strong role in resistance evolution, a small tumor-initiating population is not the only condition that restricts effective population sizes in cancer. In adjuvant therapy, a large tumor is surgically debulked before systemic treatment is initiated. Population bottlenecks created by surgical resection can considerably reduce the genetic diversity of tumor cells that escape excision. We therefore hypothesized that survival of the likeliest could play a role in the development of resistance to drugs used in the adjuvant setting.

Using a mechanistic model of tumor evolution (see Supplemental Appendix S3), we simulated tumors treated with adjuvant therapy for varying levels of pre-surgery tumor dissemination and completeness of excision. In the model, 100 initial drug-sensitive cells divide until the tumor reaches detection (10^9^-10^12^ cells), at which point the majority of the tumor is excised and the remaining population (10^5^-10^8^ cells) is treated with a small molecule inhibitor. During division events, sensitive cells can spawn one of ten resistant alleles, each with randomly preassigned allele-specific resistance and substitution likelihood parameters. For each combination of tumor size pre-and post-surgery, 50,000 patients were simulated, and their dominant resistance alleles were tallied upon relapse (i.e. when the resistant population reached a detectable size). The Spearman rank correlation between allele frequency and degree of drug resistance (Figure 6A, top) as well as allele frequency and substitution likelihood (Figure 6A, bottom) were calculated for each combination of population size pre-and post-surgery. The simulation results show that when 10^7^ or fewer cancer cells remained following resection, the allele frequency at relapse was more highly correlated with substitution likelihood than the degree of drug resistance conferred (Figure 6B).

**Figure 6:**
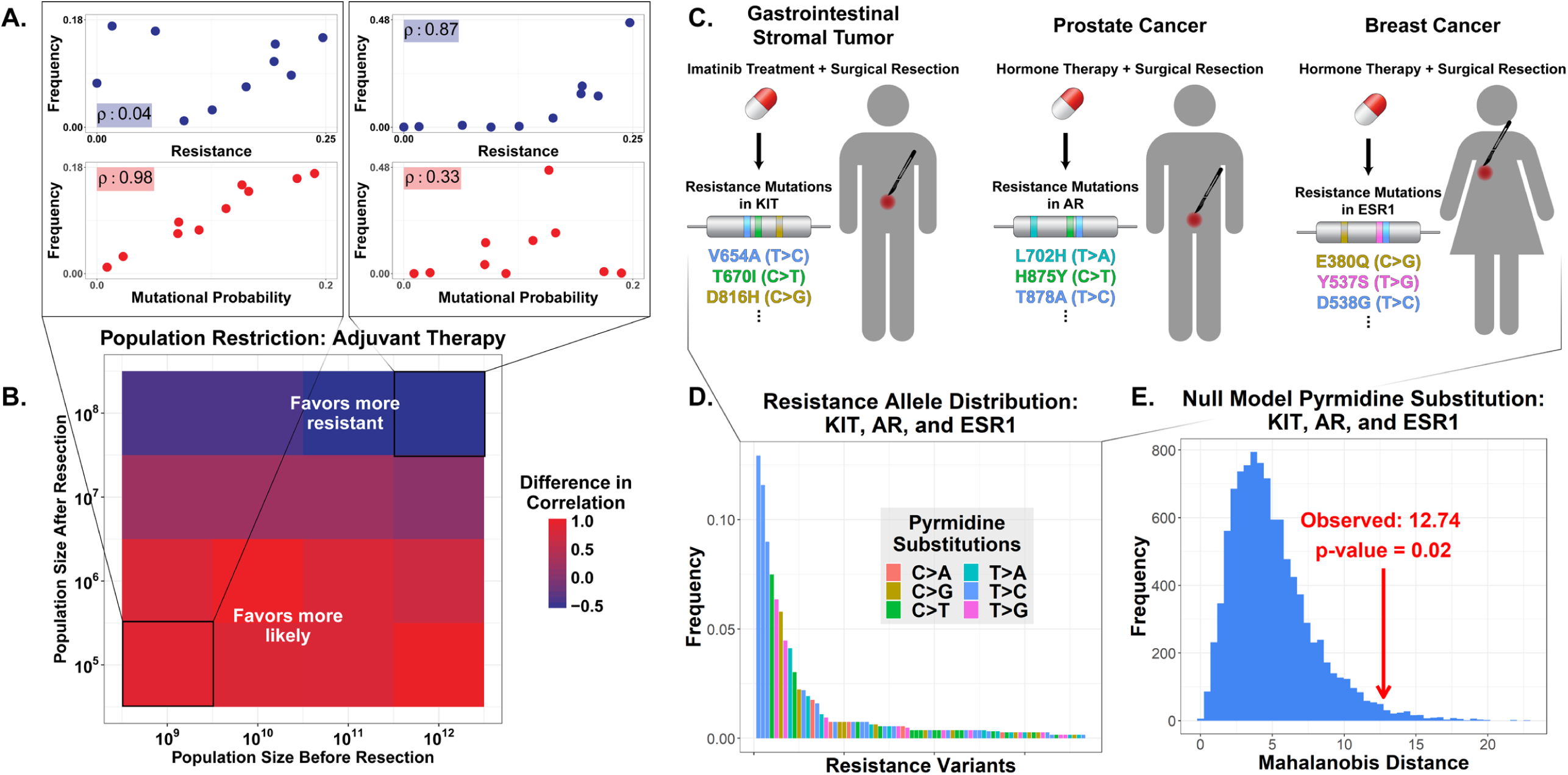
Restricted genetic heterogeneity in adjuvant therapy gives rise to survival of the likeliest. **(A)** Adjuvant therapy evolutionary model example results. Simulation results for population before resection M_pre_=10^9^ and after resection M_post_=10^5^ (*left*) and for M_pre_=10^12^ and M_post_=10^8^ (*right*). Points represent specific resistance variants and ρ values are the Spearman rank correlation between frequency and degree of resistance (*top*) and frequency and substitution likelihood (*bottom*). **(B)** Summary of simulation results for various values of M_pre_ and M_post_. Colors represent the difference in correlation (substitution likelihood ρ –resistance ρ) for each set of parameters. **(C)** Schematic detailing clinical meta-analysis. For drug targets in GIST, prostate cancer, and breast cancer, acquired resistance mutations were tallied and classified by pyrmidine substitution. **(D)** Observed resistance allele distribution for the three cancer types. Colors indicate the pyrmidine substitution associated with each mutation. **(E)** The distribution in Figure 6D was simulated by reassigning the mutation class of each variant. Each simulations’ Mahalanobis distance from the distribution of all simulations was calculated. The histogram shows the distribution of those distances. The red arrow indicates the Mahalanobis distance of the observed data from the simulated null distribution (distance = 12.74). The empirical p-value of the real-world data is 0.02, suggesting that the observed data cannot be explained by a null model with no differences in substitution likelihood.

To evaluate our theoretical results, we turned to clinical data for indications where adjuvant therapy is often used. In particular, we analyzed on-target resistance mutations in c-KIT for gastrointestinal stromal tumor (GIST) patients treated with imatinib, in ESR1 for breast cancer patients treated with ER antagonists, and in AR for prostate cancer patients treated with AR antagonists (Figure 6C). Some characteristics of the available clinical data complicate the analysis. Resistance data for GIST, breast cancer, and prostate cancer were generally not delimited by whether the drug was administered adjuvantly. This ambiguity gives rise to a less pure, more conservative data set, since our meta-analysis likely includes some late stage patients with advanced metastatic disease. This uncertainty, combined with the fact that drug fitness data is unavailable for most resistance variants in the three drug targets, precludes an analysis identical to the one taken in CML described above.

We instead asked whether the variation in resistance allele frequency across GIST, breast cancer, and prostate cancer could be explained under the null hypothesis that mutation bias is uncorrelated with clinical abundance. Previous studies have developed null models useful for detecting transition-transversion biases, even in cases of adaptive evolution^24,52^. We expanded this approach to develop a more complete model of mutational biases by considering the six possible pyrmidine substitutions (C>A, C>G, C>T, T>A, T>C, and T>G, here after referred to as mutation classes). First, we determined the distribution of resistance variants for c-KIT, ESR1, and AR for GIST, breast cancer, and prostate cancer, respectively, and noted the mutation class associated with each resistance-conferring mutation (Figure 6D and Data S2). We then simulated the same distribution under the null assumption that differences in the probabilities of mutation class do not affect resistance allele prevalence (see Supplemental Figure S4A/B and Methods). That is, we randomly reassigned the mutation class associated with each variant and then tallied the number of patients in each mutation class. Under these null simulations, all variation in resistance allele frequency must be explained by factors other than mutation bias, e.g. fitness effects, since substitution likelihood is artificially decoupled from resistance frequency. If mutation bias plays no role in the real-world allele frequency, then our observed allele frequency distribution should resemble those generated by this null model. We calculated the Mahalanobis distance from the mean for both the simulated and the observed data (Supplemental Figure S4C) and found that the observed counts of mutation class were more extreme than those simulated under the null model (empirical p-value 0.02, Figure 6E).

To validate this approach, we similarly evaluated two other data sets. The first was the resistance mutation data from our CML meta-analysis, where we anticipated a significant result implicating the role of mutation bias in allele frequency. Indeed, our analysis found an empirical p-value of 0.018 (Supplemental Figure S4E/F). The second was resistance mutation data for EGFR+ non-small cell lung carcinomas (NSCLCs) treated with erlotinib or gefitinib. There is little reason to suspect that NSCLC would have a limited effective population size, and resistance is dominated by T790M. Thus, NSCLC serves as a negative control. As expected, the observed resistance variant distribution for EGFR does not significantly differ from those generated under the null model (empirical p-value 0.38, Supplemental Figure S4G/H). These results indicate that the approach outlined here is a sensitive and specific method to identify cases where substitution likelihood shapes drug resistance evolution, and where next generation drug design could be informed by analysis of mutation biases and substitution likelihood.

## Discussion

In this study, we demonstrate the ability to accurately predict real-world resistance evolution with amino acid scale resolution. In cases of restricted genetic diversity, as in populations with small effective sizes, this predictive power requires understanding of the mutational pathways and substitution probabilities that generate the variation upon which selection acts. To be clear, it is inaccurate to say that drug selection is unimportant. Certainly, a highly likely variant that grows out during treatment must still harbor a drug resistance phenotype, and our analysis of CML epidemiological data (Figure 3) indicates that the degree of resistance conferred by a mutation is still a significant predictor of mutation prevalence. However, in CML, the ability to forecast amino acid prevalence requires an understanding of the stochastic molecular events that generate the evolution of resistance. Beyond CML, our theoretical and empirical analyses (Figure 6) indicate that drugs used in the adjuvant setting provide multiple additional examples where resistance evolution can be shaped by substitution likelihood.

Beyond the retrospective analyses of molecular evolution, our study highlights the ability to prospectively forecast macroscopic evolutionary outcomes. Recent studies have shown the power of multiscale modeling in forecasting the evolutionary trajectories of influenza evolving immune escape^17^ and predicting fitness landscapes of bacteria evolving resistance against trimethroprim^26^. Our work builds on these findings by predicting epidemiology-scale resistance from molecular insights. While *in vitro* parallel evolution experiments have been able to qualitatively nominate resistance variants (Supplemental Figure S5), ours is the first study to quantitatively predict interpatient heterogeneity with a completely mechanistic model of intratumoral heterogeneity. Moreover, we concretely demonstrate how these models could be used to optimize rational drug design.

Despite their power, our predictions are an imperfect step forward. There is unfit variance in our statistical model of the epidemiological data (the predictions from our mechanistic model has a correlation of 0.86 with the clinical data). While the accuracy of these models is surprisingly high given their simplicity and the fact that none of the mechanistic parameters were fit to clinical data, there are other variables that could explain the small amount of residual variance. The first is that the genetic background outside of ABL1 can alter the level of phenotypic drug resistance. The second is that *in vivo* niches can provide protection from drug effects and/or exposure ^53,54^, and if the fitness as determined by the niche is variable across patients or mutations, this could strongly contribute to existing uncertainty in our predictions. Finally, it is also possible that non-genetic single cell heterogeneity contributes to the residual error in the model^55^. Yet despite these potential sources of unmeasured error, we explain most of the clinical phenotypes that we aimed to predict.

The basic concept of evolutionarily informed treatment regimens that minimize drug resistance is not new. Theoretical evolutionary models^10,11^ support empirical clinical evidence^56^ that non-cross-resistant combination therapy can result in durable responses for some patients. However, effective non-cross-resistant drug combinations are not available for many cancers and multiple drugs can cause overlapping off-target toxicities. To our knowledge, our study is the first to identify a quantitative evolutionary design principle for single agent therapy. When effective population sizes are small, evolution favors the most likely resistance mutation and so should drug discovery (Figure 5). This is not simply a matter of trying to reduce the absolute number of mutational paths to drug resistance. Prospective therapeutic leads can be prioritized for evolutionary optimality even when they have an identical absolute number of mutational liabilities (as in Figure 5A). Moreover, even when a combination can be identified, treatment non-adherence and other resistance risks would suggest that optimal single agents should enable combinations that are less evolutionarily risky. This would suggest that optimized single agents could help create optimized drug combinations.

Predictive evolutionary models could also inform the design of frontline drugs with second-and third-generation drug liabilities in mind. Frontline therapies select for resistance variants that eventually determine the dominant tumor genotype upon treatment failure. The patient is then treated with next-generation drugs, if any exist, until they too fail. Every iteration of this process determines the genetic background that the next drug will face, as each treatment selects for subclones among the population of cells resistant to the treatment before it. Moreover, next generation drug discovery is currently entirely empirical as physicians and drug discovery scientists simply take the next step based upon the liabilities of their last step. Cancer offers a unique opportunity to escape this arms race because every new patient presents with drug naïve disease. Predictive modeling could be used to design drugs that will select for the most favorable genotype for the next line of treatment, and allow clinicians to more successfully shape the broader evolutionary trajectory of a tumor. Thus our discoveries could enable concepts like collateral sensitivity^57^ by providing a model-driven framework.

In diseases with a high degree of heterogeneity, even one resistance liability is sufficient to virtually guarantee treatment failure. This is not true when genetic diversity is limited by the effective population size. In this study, we’ve predicted that targeting the most likely clinical mutations during drug design is a potential strategy to minimize the prevalence of resistance across a population. This raises the possibility that targeted therapies that are used in the adjuvant setting may be designed differently than therapies developed to treat large metastatic disease burden. In the future, we believe that rational drug design will be synonymous with evolutionarily-informed drug design. Predictive evolutionary models will enable precision medicines that are more efficacious and longer lasting in the face of biological change.

## Methods

### Construct generation

Following recombination-based cloning into pLVX-IRES-Puro, site directed mutagenesis was utilized to make the correct mutation in BCR-ABL. Mutation identity was confirmed by Sanger sequencing.

### Cell line generation

BaF3 cells were ordered from DSMZ. BaF3 cells are maintained in 1640 (Sigma Aldrich) +10%FBS(Fisher)+1%Penn/Strep (Life Technologies) and 10ng/mL IL-3 (PeproTech). Lentiviral constructs were co-transfected with calcium phosphate alongside third generation packaging vectors that were pseudotyped with VSV-G. Viral supernatant was collected at 24 hours^58^. All BCR-ABL mutations were infected at limiting MOI to achieve the lowest viral titer required to produce IL-3 independence. After selection in the absence of IL-3, we tested for puromycin resistance. An assumption of BaF3 usage is that the sensitivity or resistance seen in a BaF3 cell is a reasonable approximation of the resistance and sensitivity seen in a human leukemia. This assumption has been shown to be a reasonable approximation in numerous prior studies^30,33^. Moreover, the number and diversity of mutants that were explored in this study could not be achieved by using primary samples. The rarity of CML and the limited patients observed make the collection of primary tissue harboring all mutations essentially impossible. All engineered cell lines were sequenced in the BCR-ABL kinase domain to confirm their identity.

### IC_50_ measurements

Eleven serial dilutions were performed. Imatinib, dasatinib and nilotinib were all obtained from Selleck Chem. Starting points for dilutions were 10uM, 100nM, and 1uM respectively. 3000 BaF3 cells with the indicated mutation were seeded into a 96 well plate in 150ul of RPMI 1640 (Sigma Aldrich) +10%FBS (Fisher)+1%Penn/Strep (Life technologies). After addition of the drugs, cells were left in the incubator for 72 hours. At 72 hours Cell Titer Glo (Promega) was added at 1:4. This is less than the manufacturer’s instructions, because we have verified the sensitivity of the assay with this reduced protocol. Plates were read via a luminescence plate reader at 72 hours.

### Serum protein shifts

HSA-AAG containing medium is RPMI 1640 growth medium with the addition of 341 mM HSA (human serum albumin, Sigma, Cat # A9511) and 1 mg/ml AAG (human a1-acid glycoprotein, Sigma, Cat # G9885). All medium is sterilized by filtration through a 0.22mm membrane. Following the formulation of HSA-AAG media, identical IC_50_ curve experiments are performed. The fold change of the IC_50_ across 3 mutants of varying affinities (M244V, E255V, and WT BCR-ABL were used). Serum protein binding decreases the free drug concentration and shifts IC50 values to numbers that are directly comparable to measured *in vivo* pharmaceutical exposures.

### Exome data analysis

Data from the broad ExAC consortium (http://exac.broadinstitute.org/) for ABL1 was downloaded as of 11-17-2017. Raw data and processing code are available in Figure 3 in our GitHub repository: https://github.com/pritchardlabatpsu/PredictiveResistanceEvolution/tree/master/Figures/Figure3/Mutation%20probabilities We tallied each of the 12 possible nucleic acid substitutions (not six) because transcription coupled repair has been shown to cause biases in the mutational spectrum of the transcribed strand, and ABL1 is widely expressed. We also compared the distribution of substitutions across ABL1 in the ExAC data to genome wide mutation biases measured in CML exomes, as well as simple transition/transversion biases from the literature, and we did not observe significant differences.

### Clinical data

Clinical data was identified from six studies and the Wellcome Trust. The Sanger Wellcome Trust download was as of 12-01-2017. (https://cancer.sanger.ac.uk/cosmic/csamples/details).

### Theoretical Model of Competing Resistance Alleles

See Appendix S1 and code repository. .m files are included under the Figure 2C folder on GitHub. https://github.com/pritchardlabatpsu/PredictiveResistanceEvolution/tree/master/Figures/Figure2

### Epidemiological Analysis

See Supplemental R markdown files and code repository for Figure 3. Files were written in R markdown. Raw data and analysis code (.Rmd and knitted .html files) are provided on GitHub: https://github.com/pritchardlabatpsu/PredictiveResistanceEvolution/tree/master/Figures/Figure3

### CML Model

See Appendix S2 and code repository. All raw input data, source code, and simulation outputs are available on GitHub for Figures 4. .m files, simulation outputs, and .Rmd files for imatinib, nilotinib, and dasatinib are available in the Figure 4 folder: https://github.com/pritchardlabatpsu/PredictiveResistanceEvolution/tree/master/Figures/Figure4 The analysis of maxitinib in figure 5 can be found at GitHub: https://github.com/pritchardlabatpsu/PredictiveResistanceEvolution/tree/master/Figures/Figure5

### Adjuvant Therapy Model

See Appendix S3 and code repository. All code, parameters, outputs, and a LaTeX file describing the models are available at https://github.com/pritchardlabatpsu/PredictiveResistanceEvolution/tree/master/Figures/Figure6

### Clinical adjuvant therapy analysis

We compiled count data for resistance variants in c-KIT, ESR1, and AR for GIST, breast cancer, and prostate cancer, respectively, and classified each by the associated pyrmidine substitution (C>A, C>G, C>T, T>A, T>C, and T>G; mutation class). For cases when an amino substitution could be caused by multiple single nucleotide mutation paths, we classified them when nucleotide substitution resolution was available in the literature and excluded the counts otherwise. Under the null model, these variants were randomly reassigned mutation classes and the sum of all counts for each mutation class was noted. Since our data only considers mutations that result in an amino acid change, the mutation class assignments were weighted by the number of mutations in each class that are nonsynonymous (68 C>A; 76 C>G; 58 C>T; 64 T>A; 59 T>C; 68 T>G). 10,000 replicates were simulated for each data set. The Mahalanobis distance of each simulation from the mean of all simulations was calculated; the Mahalanobis distance of the observed mutation class counts from the simulation mean was also determined. An empirical p-value was obtained as *p* = *r*/*n*, where *n* is the number of simulations and *r* is the number of simulations with a Mahalanobis distance greater than or equal to that of the observed data. Identical approaches were taken for resistance variant counts for ABL1 in CML and EGFR in NSCLC. Raw data and processing code are available in our GitHub repository: https://github.com/pritchardlabatpsu/PredictiveResistanceEvolution/tree/master/Figures/Figure6

## Data and Code Availability

https://github.com/pritchardlabatpsu/PredictiveResistanceEvolution/

## Acknowledgements

The authors would like to thank Vivek Kapur, Mike Schmitt and Victor Rivera for important conceptual and technical discussions that heavily influenced the direction of the manuscript. Mengrou Lu, Kyle McIlroy, Lauren Randolph, Maciej Boni, Jeremy Rock, and David Kennedy provided important contributions to the manuscript through critical comments, intellectual insights, or writing/figure assistance. Igor Aronson provided critical assistance in the mathematics underlying Figure 2. Members of the PSU Resistance Cluster, and attendees at the 2018 Gordon conference on drug resistance were also critical to shaping this manuscript.

## Notes

https://github.com/pritchardlabatpsu/PredictiveResistanceEvolution

