## Supplemental Figures for "Multi-scale predictions of drug resistance epidemiology suggest a new design principle for rational drug design"

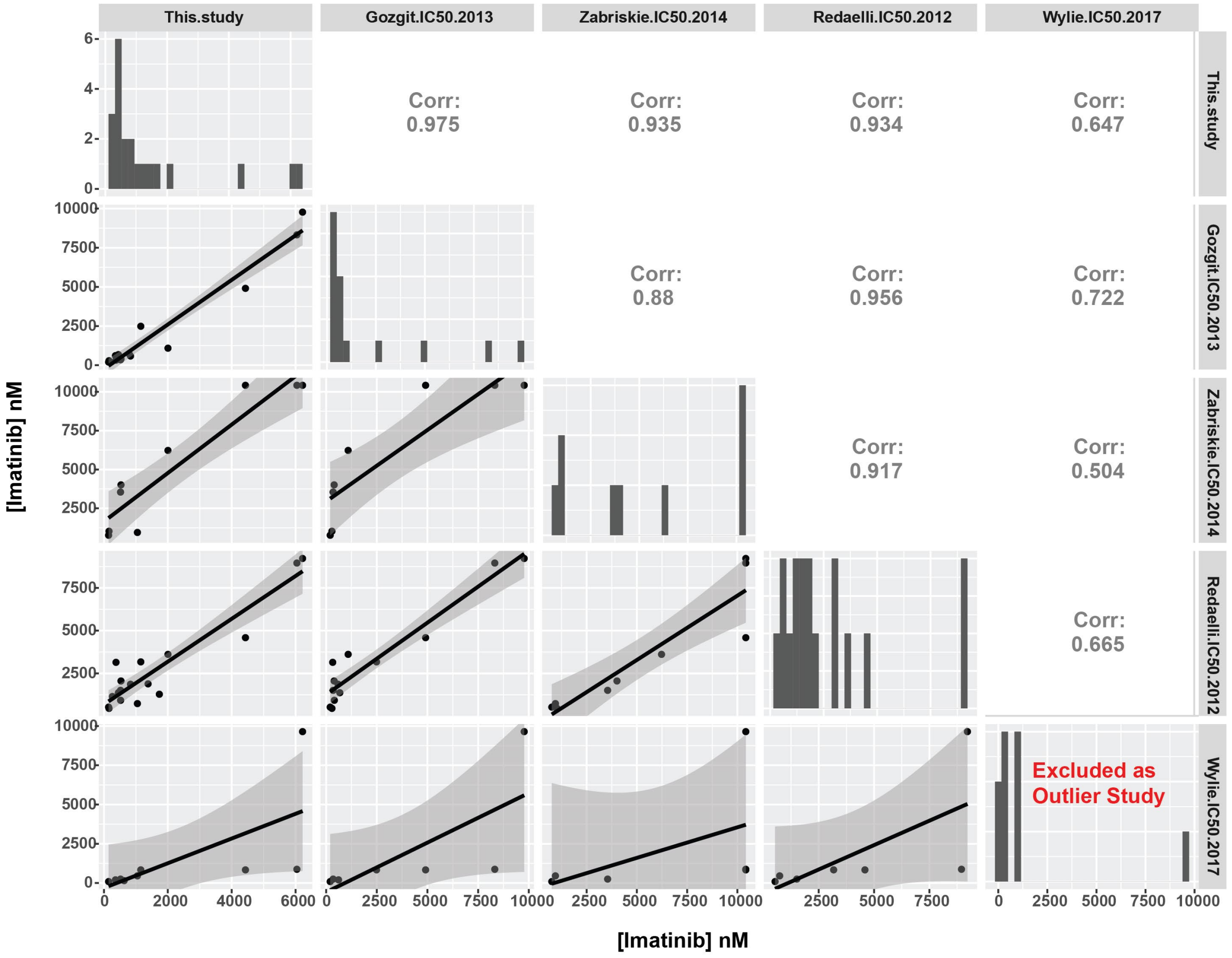


**Supplemental Figure S1: Analysis of $IC_{50}$ measurements in BCR-ABL resistance mutants across studies.**

We analyzed the $IC_{50}$ of BaF3s transduced with BCR-ABL variants as reported in other studies. The diagonal of the figure shows the histogram of $IC_{50}$s for each study. The correlations are reported in the off-diagonals. In general, four studies (ours, Gozgit 2013, Zabriskie 2014, and Redaelli 2012) exhibit high cross-correlation (Pearson’s rho ≥ 0.88). The Wylie 2017 study was excluded from further analysis given its low correlation with the remaining four studies. See Methods and Table S1 for explanation of the normalization procedure.

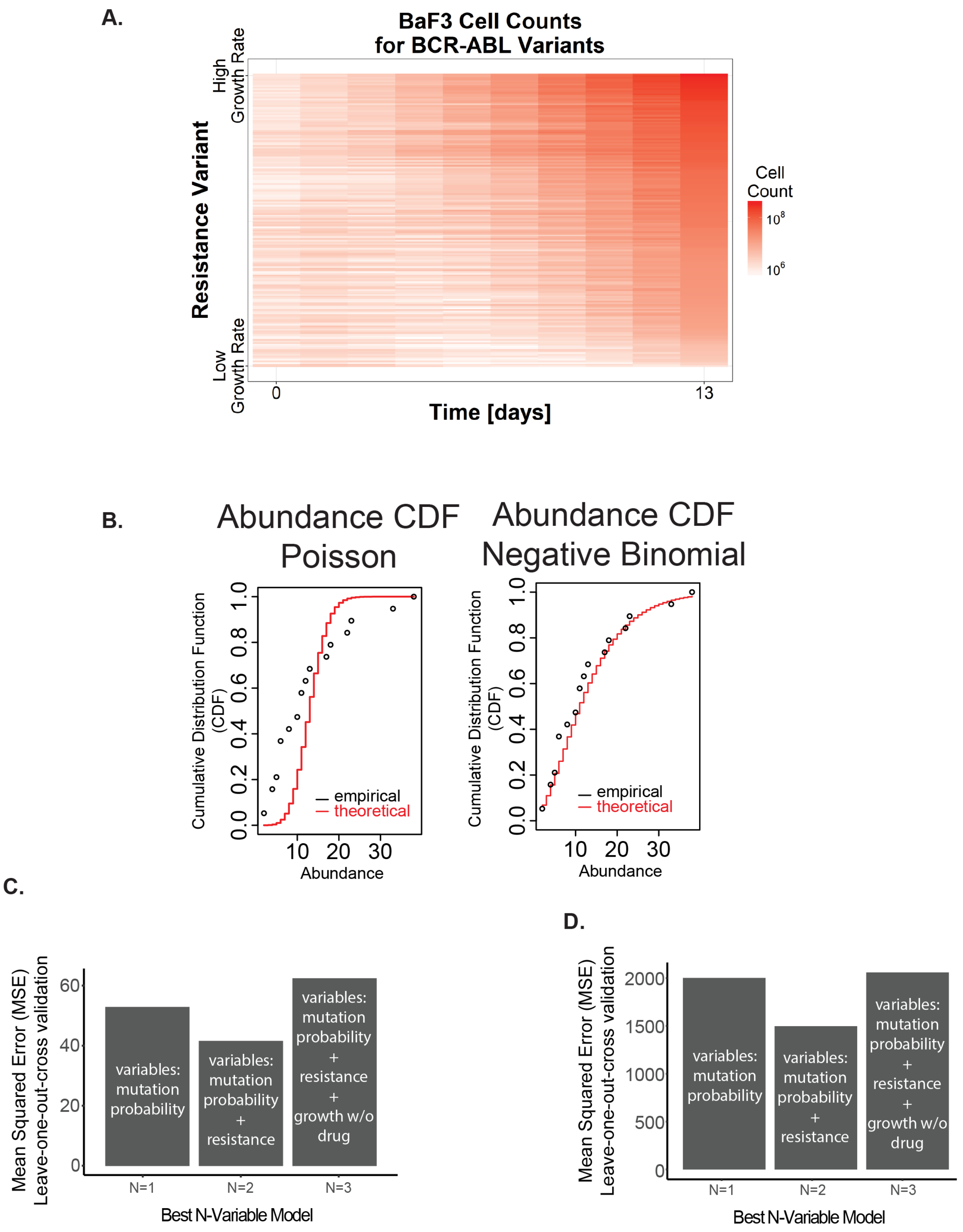
A.
BaF3 Cell Counts
for BCR-ABL Variants
Resistance Variant
High
Growth Rate
Low
Growth Rate
Time [days]
0
13
Cell
Count
$10^8$
$10^6$
B.
Abundance CDF
Poisson
Abundance CDF
Negative Binomial
Cumulative Distribution Function
(CDF)
0.0
0.2
0.4
0.6
0.8
1.0
10
20
30
Abundance
empirical
theoretical
C.
Mean Squared Error (MSE)
Leave-one-out-cross validation
0
20
40
60
variables:
mutation
probability
variables:
mutation
probability
+
resistance
variables:
mutation
probability
+
resistance
+
growth w/o
drug
N=1
N=2
N=3
Best N-Variable Model
D.
Mean Squared Error (MSE)
Leave-one-out-cross validation
0
500
1000
1500
2000
variables:
mutation
probability
variables:
mutation
probability
+
resistance
variables:
mutation
probability
+
resistance
+
growth w/o
drug
N=1
N=2
N=3
Best N-Variable Model

**Supplemental Figure S2: Construction of resistance mutation prevalence model.**
A statistical model was developed to explain the variance in clinical prevalence of BCR-ABL resistance mutations. **(A)** To construct the model, we considered three potential predictor variables: imatinib $IC_{50}$ (Figure 3B), growth rate in the absence of drug (Supplemental Figure 2A), and amino acid substitution probability (Figure 3D). **(B)** The amino acid prevalence assembled from our meta-analysis of multiple clinical trials was analyzed to identify the appropriate generalized linear model to use for regression. The empirical CDF of the prevalence count data was compared to theoretical CDFs given a Poisson (left) and negative binomial distribution (right). The Poisson distribution was overdispersed, while the negative binomial distribution appeared to agree with the empirical data and so a negative binomial regression model was used. **(C)** Validation of the regression model trained on our clinical meta-analysis data set. Leave-one-out cross validation was used to estimate the test error of the N-variable model with the lowest AIC for each N. The N=1 model includes substitution bias; the N=2 model includes substitution bias and $IC_{50}$; the N=3 model includes substitution bias, $IC_{50}$, and growth rate in the absence of drug. The N=2 model had the lowest estimated mean square error. For a complete explanation of model construction see the Figure 3 R-markdown file on GitHub. **(D)** Validation of regression model as in **(C)** for model trained on the Sanger Institute data set.

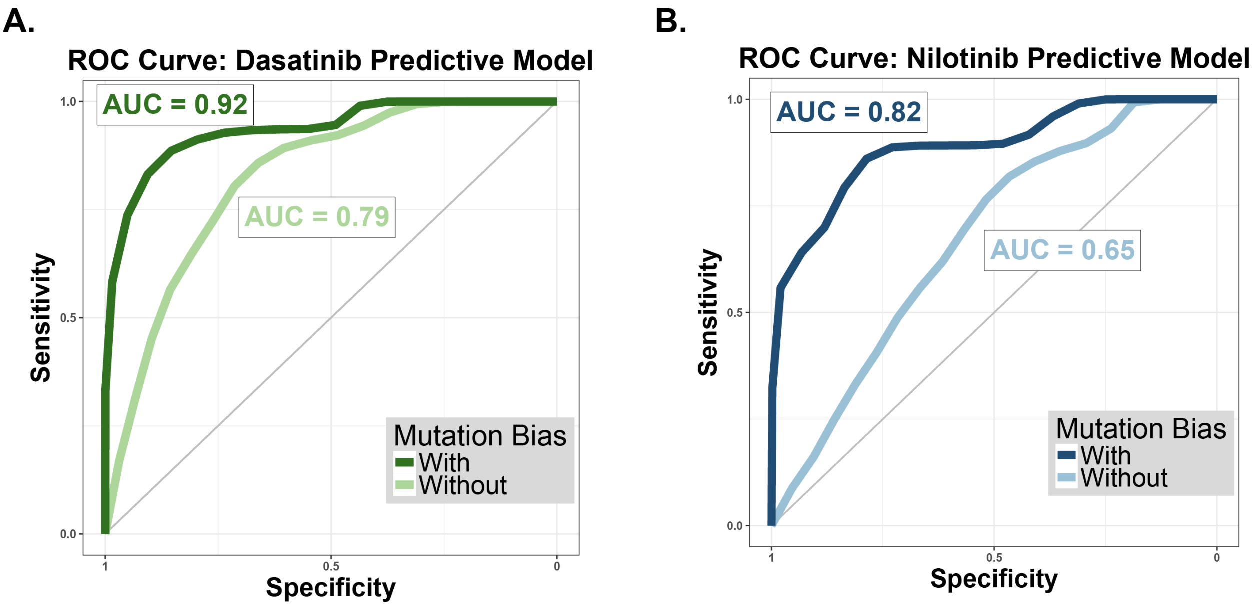


**Supplemental Figure S3: Predictive modeling can predict which resistance mutants are present in a clinical trial**

**(A)** ROC curve for dasatinib predictive model. The results of our dasatinib simulations were downsampled to reflect the size of frontline dasatinib trial. For each downsampling, a binary classifier was built from the predicted allele frequency, and an ROC curve was constructed. The mean ROC curve across simulations for a model with (**dark green**) and without (**light green**) mutation bias is shown here. AUC represents area under ROC curve. **(B)** Mean ROC curve for nilotinib predictive model, with (**dark blue**) and without (**light blue**) mutation bias.

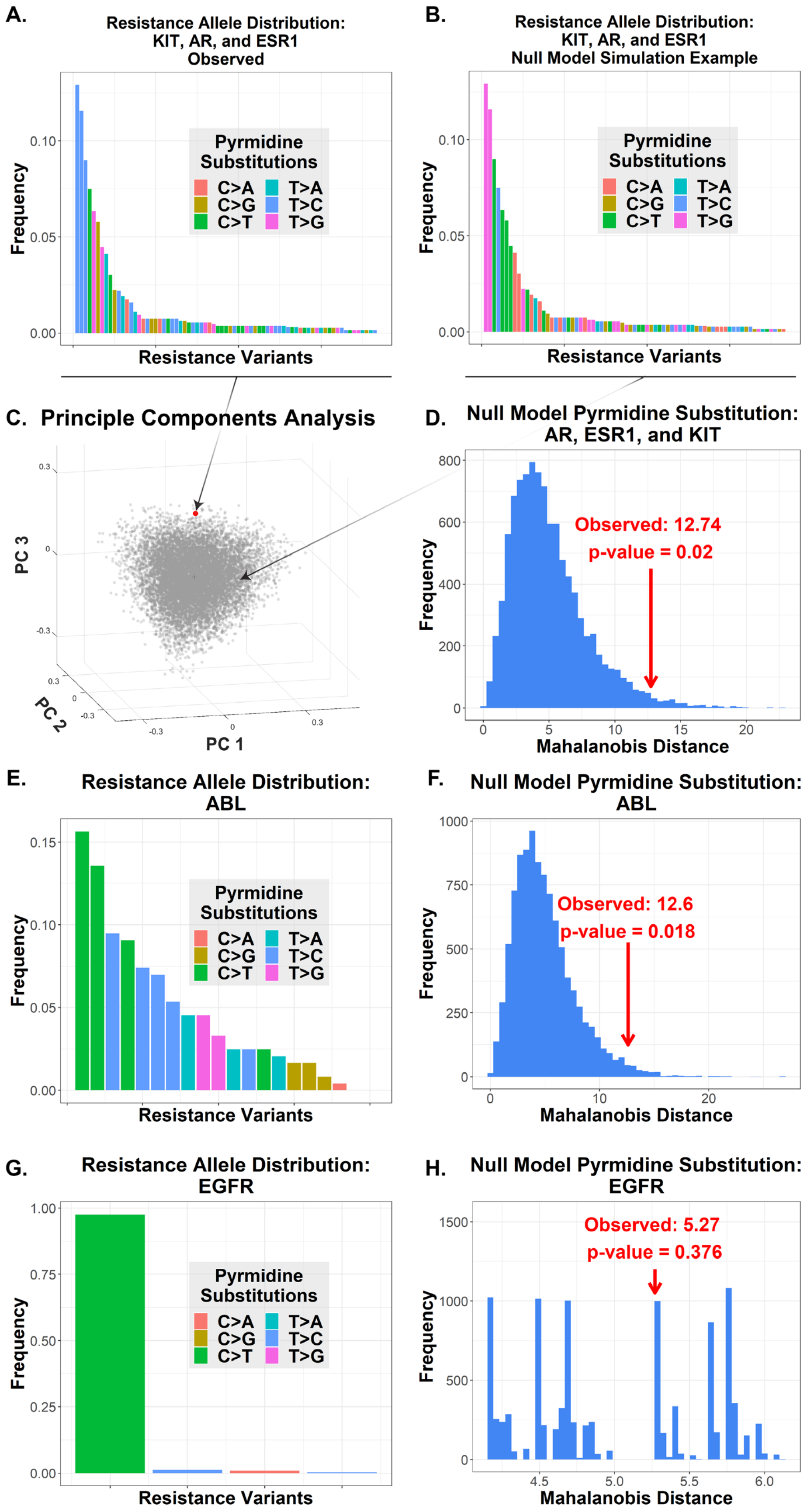
A.
Resistance Allele Distribution:
KIT, AR, and ESR1
Observed
Pyrmidine Substitutions
C>A C>G C>T T>A T>C T>G
Frequency
Resistance Variants
B.
Resistance Allele Distribution:
KIT, AR, and ESR1
Null Model Simulation Example
Resistance Variants
C. Principle Components Analysis
PC 1
PC 2
PC 3
D. Null Model Pyrmidine Substitution:
AR, ESR1, and KIT
Observed: 12.74
p-value = 0.02
Mahalanobis Distance
E. Resistance Allele Distribution:
ABL
F. Null Model Pyrmidine Substitution:
ABL
Observed: 12.6
p-value = 0.018
G. Resistance Allele Distribution:
EGFR
H. Null Model Pyrmidine Substitution:
EGFR
Observed: 5.27
p-value = 0.376

**Supplemental Figure S4: Resistance to drugs used in the adjuvant setting exhibit survival of the likeliest**

**(A)** The observed distribution of resistance mutations in KIT, AR, and ESR1 for GIST, prostate cancer, and breast cancer patients, respectively. Colors indicate mutation class, i.e. pyrmidine substitution. **(B)** A sample simulation of the KIT/AR/ESR1 resistance allele distribution generated under a null model. The frequency distribution is preserved across simulations, but each variant is randomly reassigned a mutation class. For each simulation, the sum total of patients associated with each mutation class is noted. **(C)** A projection of the simulation results (**gray**) along the first three principle components. The observed data from our clinical meta-analysis is shown in **red**. **(D)** The Mahalanobis distance of each simulation from the distribution of all simulations was calculated. The distribution of Mahalanobis distances is shown here. The red arrow indicates the Mahalanobis distance of the observed data from the simulated distribution (empirical p-value 0.02). **(E)** An identical approach was used to analyze the CML data, which served as a positive control. Shown here is the distribution of ABL1 resistance alleles categorized by mutation class. **(F)** The distribution of Mahalanobis distances for the randomly reassigned CML data. The distance of the observed data from the null distribution is 12.6 (p-value = 0.018). **(G)** The same approach was used to analyze resistance prevalence data for EGFR inhibitors in NSCLC, which served as a negative control. The distribution of resistance alleles is shown. **(H)** The distribution of Mahalanobis distances for the simulated EGFR data. Observed distance is 5.27 (p-value = 0.376), indicating a nonsignificant result as expected.

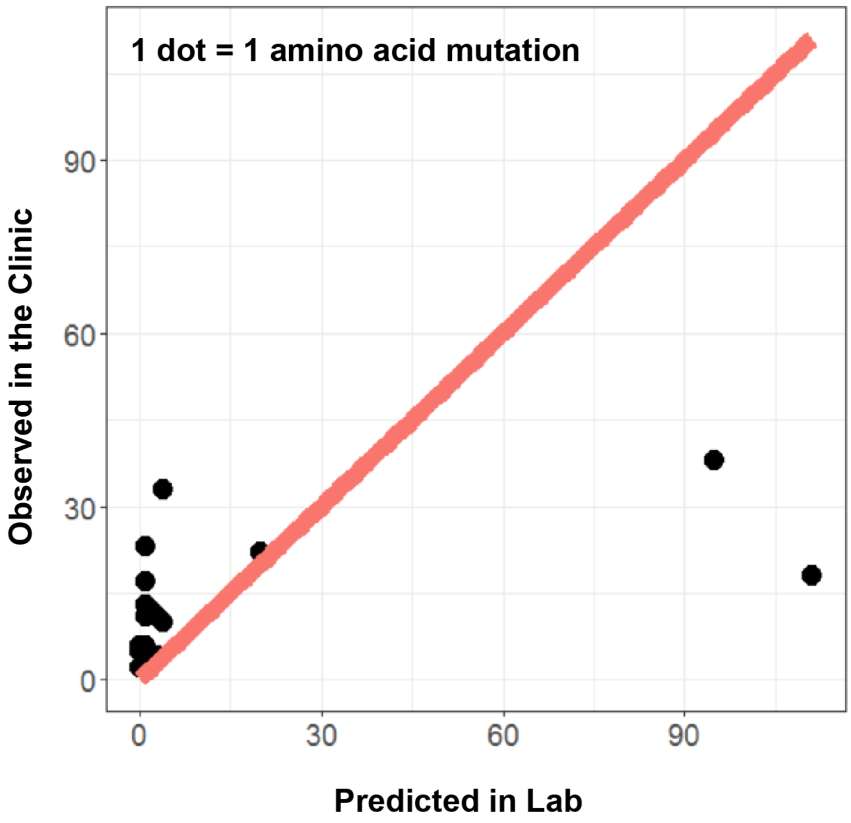


**Supplemental Figure S5: ENU mutagenesis qualitatively identifies resistance mutations but fails to quantitatively predict abundance.**
Bradeen et al. have previously used ENU mutagenesis to nominate imatinib resistance mutations in BCR-ABL. The abundance of each mutation as predicted by the observation frequency in the mutagenesis experiment is given on the x-axis. The observed abundance of each mutation from our meta-analysis of clinical trials is given on the y-axis. The results indicate some qualitative concordance of resistance phenotypes between mutants predicted in the lab and those observed clinically. However, the *in vitro* mutagenesis experiment poorly predicts the quantitative frequency of mutations observed *in vivo*.
