## Supplemental Appendix 1 for "Multi-scale predictions of drug resistance epidemiology suggest a new design principle for rational drug design"

### Supplemental Appendix S1:
#### Analytical Two-Allele Probability Model

Leighow and Liu et al.

July 14, 2019

### 1 Introduction

Consider a population of pathogenic, asexual cells which expand prior to drug treatment and collapse during treatment. During the lifetime of this population of cells, the initial sensitive population has the potential to seed resistant subclones which have a positive net growth rate in the presence of drug. The population dynamics of a resistant clone depend on allele-specific parameters, in particular its mutational probability (i.e. the likelihood that the resistant clone will spawn) and its degree of resistance (i.e. the growth rate during treatment).

We are interested in identifying the conditions for which mutation bias plays a significant role in determining the prevalence of a given mutation at the scale of patient populations. To address this, we consider the simplest case of two distinct mutations that confer drug resistance (Allele A and Allele B). Given their rarity and to simplify the analysis, we do not consider the potential for compound mutations.

#### 1.1 Resistance and Mutational Probability Profiles

It is interesting to consider the case where there exists a trade-off between resistance and mutational probability. We assign Allele A a high degree of resistance and low mutational probability; we assign the opposite profile for Allele B. These profiles are summarized in **Appendix Table 1.1**.

**Appendix Table 1.1**

| | Allele A | Allele B |
|---|---|---|
| Resistance | high | low |
| Mutational Probability | low | high |

We then ask the question: which allele will drive relapse? In other words, which is the first clone to reach clinical detection? In identifying conditions where Allele B (the allele with a greater mutational probability but a lower degree of resistance) is more likely to be the dominant allele, we find parameter sets where we expect mutational probability to play a significant role in shaping distributions of specific amino acid mutations in clinical populations.

#### 2 General Outcomes

Towards identifying which allele will drive relapse in a two-allele system, consider the four possible outcomes:

**Appendix Table 1.2**

| | | Allele A present? | |
|---|---|---|---|
| | | No | Yes |
| Allele B present? | No | **Outcome 1** | **Outcome 2** |
| | Yes | **Outcome 3** | **Outcome 4** |

Here, we say an allele is present if its mutational event occurs during the lifetime of the sensitive population and the newly spawned clone escapes stochastic eradication (i.e. it eventually reaches clinical detection). Thus, the results of each outcome are summarized as follows.

**Outcome 1**: Neither allele is present.

**Outcome 2**: Allele A is fixed.

**Outcome 3**: Allele B is fixed.

**Outcome 4**: Both alleles are present.

**Outcome 4** can be further divided into two suboutcomes:

**Outcome 4.A**: Allele A reaches clinical detection first.

**Outcome 4.B**: Allele B reaches clinical detection first.

Towards addressing our initial question, we see that:

$$\begin{aligned} &P[\text{Allele A is dominant upon relapse}] = \\ &\qquad P[\textbf{Outcome 2}] + P[\textbf{Outcome 4.A}|\textbf{Outcome 4}] \cdot P[\textbf{Outcome 4}] \\ &P[\text{Allele B is dominant upon relapse}] = \\ &\qquad P[\textbf{Outcome 3}] + P[\textbf{Outcome 4.B}|\textbf{Outcome 4}] \cdot P[\textbf{Outcome 4}] \end{aligned} \tag{1}$$

In the following analyses, we will derive these probabilities.

#### 3 Probability of Mutation

Let us begin by considering the probabilities of each outcome. Because the presence of each allele is independent of the other, it follows that:

$$
\begin{aligned}
P[\textbf{Outcome 1}] &= P[\text{Allele A is absent}] \cdot P[\text{Allele B is absent}] \\
P[\textbf{Outcome 2}] &= P[\text{Allele A is present}] \cdot P[\text{Allele B is absent}] \\
P[\textbf{Outcome 3}] &= P[\text{Allele A is absent}] \cdot P[\text{Allele B is present}] \\
P[\textbf{Outcome 4}] &= P[\text{Allele A is present}] \cdot P[\text{Allele B is present}]
\end{aligned} \tag{2}
$$

To find the probability that an allele is absent, let $N_i(t)$ be the number of mutational events generating the $i$th allele that occur on the time interval $(0, t]$. The discrete random variable $N_i(t)$ can be modeled as a nonhomogenous Poisson process. Thus, the probability that $k$ mutations occur on the interval $(0, t]$ is given by:

$$
P[N_i(t) = k] = \frac{[\int_0^t \lambda_i(y)dy]^k}{k!} \exp\left( - \int_0^t \lambda_i(y)dy \right) \tag{3}
$$

where the hazard rate $\lambda_i(t)$ is the probability of a mutation to the $i$th allele occurring on the infinitesimal interval $(t, t + dt)$. This hazard rate is a function of the size of the sensitive population $x_S(t)$ from which the resistant variant arises, and is given by:

$$
\lambda_i(t) = x_S(t) b_S \mu \rho_i \frac{r_i}{b_i} \tag{4}
$$

Here, $b_S$ is the division rate of the sensitive population; $\mu$ is the probability that a resistance mutation occurs per division event of the sensitive population; $\rho_i$ is the probability of mutation to the $i$th allele given that a resistance mutation occurs. The parameter $\rho_i$ is used to describe differences in allele-specific mutational probabilities. Note that $\sum_i \rho_i = 1$. We multiply by the allele-specific survival probability $r_i/b_i$ (previously derived by Athreya and Ney (2004) [**1**]) to consider only mutation events where the newly spawned clone escapes stochastic eradication. The parameters $r_i$ and $b_i$ are the net growth rate and division rate of the $i$th resistant population, respectively.

The size of the sensitive population can be modeled assuming exponential growth prior to treatment and exponential decay during treatment. If treatment starts at time $T$, then the number of sensitive cells $x_S(t)$ is:

$$
x_S(t) = \begin{cases} \frac{b_S}{r_S} e^{r_S t} & 0 \le t < T \\ M e^{r'_S (t-T)} & T \le t \end{cases} \tag{5}
$$

Here, $r_S$ is the net growth rate of sensitive cells prior to treatment, $r'_S$ is their net growth rate during treatment ($r'_S < 0$) and $M$ is the size of the sensitive population at the start of treatment (i.e. the detection size).

The time to detection $T$ can be related to $M$ by solving the equation $\frac{b_S}{r_S} e^{r_S T} = M$, yielding:

$$
T = \frac{1}{r_S} \ln\left( \frac{M r_S}{b_S} \right) \tag{6}
$$

Substituting equations (4) and (5) into (3) and choosing $k = 0$ events, we arrive at an expression $S_i(t)$ for the probability that a mutation to the $i$th allele has not occurred by time $t$.

$$S_i(t) = P[N_i(t) = 0] = \begin{cases} \exp\Big( - \frac{b_S^2 r_i \mu \rho_i}{r_S b_i} \int_0^t e^{r_S y} dy\Big) & 0 \leq t < T \\ \exp\Big( - \frac{b_S^2 r_i \mu \rho_i}{r_S b_i} \int_0^T e^{r_S y} dy - \frac{M b_S r_i \mu \rho_i}{b_i'} e^{-r_S' T} \int_T^t e^{r_S' y} dy\Big) & T \leq t \end{cases} \tag{7}$$

Let $\alpha_i = \frac{b_S^2 r_i \mu \rho_i}{r_S b_i}$ and $\beta_i = \frac{M b_S r_i \mu \rho_i}{b_i'} e^{-r_S' T}$. Solving the integrals, we can simplify equation (7) to:

$$S_i(t) = \begin{cases} \exp\Big( - \frac{\alpha_i}{r_S}(e^{r_S t} - 1)\Big) & 0 \leq t < T \\ \exp\Big( - \frac{\alpha_i}{r_S}(e^{r_S T} - 1) - \frac{\beta_i}{r_S'}(e^{r_S' t} - e^{r_S' T})\Big) & T \leq t \end{cases} \tag{8}$$

If we define $\eta_i = e^{\alpha_i / r_S}$ and $\nu_i = \eta_i \exp[\frac{-\alpha_i}{r_S} e^{r_S T} + \frac{\beta_i}{r_S'} e^{r_S' T}]$, $S_i(t)$ can be further simplified:

$$S_i(t) = \begin{cases} \eta_i \exp\Big( \frac{-\alpha_i}{r_S} e^{r_S t}\Big) & 0 \leq t < T \\ \nu_i \exp\Big( \frac{-\beta_i}{r_S'} e^{r_S' t}\Big) & T \leq t \end{cases} \tag{9}$$

The complement of $S_i(t)$, $F_i(t) = 1 - S_i(t)$, gives the cumulative mutational risk, i.e. the probability that at least one mutation *has* occurred by time $t$. An example trajectory of $F_i(t)$ is shown in **Appendix Figure 1.1**.

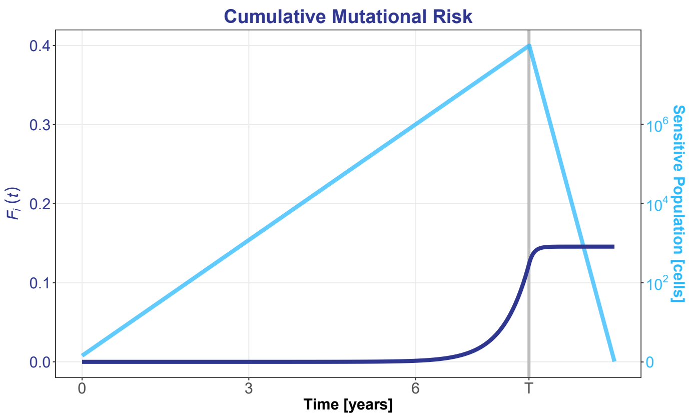


**Appendix Figure 1.1**: $F_i(t)$ for $b_S = b_i = 0.0088$ /day, $r_S = r_i = 0.0062$ /day, $r'_S = -0.0329$ /day, $r'_i = 0.0060$ /day, $\mu = 4 \times 10^{-8}$ /division, $\rho_i = 0.03$, and $M = 10^8$ cells. Growth parameters from estimates on leukemic cell dynamics reported by Fassoni et al (2018) [**2**] given a turnover rate of 0.3 (estimate from Komarova and Wodarz (2005) [**3**]). Resistance allele-specific parameters chosen to match E255V. Mutation rate from Michor et al (2005) [**4**].

Recalling that $r'_S < 0$, the probability that no mutation occurs ($S_i(\infty)$) or at least one mutation occurs ($F_i(\infty)$) during the lifetime of the tumor is given by evaluating equation (9) for $t \to \infty$:

$$\begin{aligned} S_i(\infty) &= \nu_i \\ F_i(\infty) &= 1 - S_i(\infty) = 1 - \nu_i \end{aligned} \tag{10}$$

This is a generalization of a result previously obtained by Bozic et al (2013) [**5**].

This gives us expressions for the probabilities of the four outcomes outlined in **Appendix Table 1.1**:

$$\begin{aligned} P[\textbf{Outcome 1}] &= S_A(\infty) \cdot S_B(\infty) = \nu_A \cdot \nu_B \\ P[\textbf{Outcome 2}] &= F_A(\infty) \cdot S_B(\infty) = (1 - \nu_A) \cdot \nu_B \\ P[\textbf{Outcome 3}] &= S_A(\infty) \cdot F_B(\infty) = \nu_A \cdot (1 - \nu_B) \\ P[\textbf{Outcome 4}] &= F_A(\infty) \cdot F_B(\infty) = (1 - \nu_A) \cdot (1 - \nu_B) \end{aligned} \tag{11}$$

### 4 Outcome 4: Competition

We now turn our attention to **Outcome 4**, where both alleles are present. Since we are interested in identifying which allele drives relapse, we ask the question: which clone will reach clinical detection first? The first resistant subclone to reach clinical detection determines the time scale for relapse, retreatment, and patient survival, and thus has greater immediate clinical significance than a minor subclone. Here, we assume clinical detection of the resistant clone occurs at the same size as for the primary sensitive tumor, i.e. $M$.

We see that

$$\begin{aligned} P[\textbf{Outcome 4.A}] &= P[T_{det,A} < T_{det,B}] \\ P[\textbf{Outcome 4.B}] &= P[T_{det,B} < T_{det,A}] \end{aligned} \tag{12}$$

where $T_{det,i}$ is the time to detection of the $i$th allele (i.e. the time for the $i$th allele to reach detection size $M$). We focus on finding an expression for the probability of **Outcome 4.B**, since the probability of **Outcome 4.A** is easily found using $P[\textbf{Outcome 4.A}|\textbf{Outcome 4}] = 1 - P[\textbf{Outcome 4.B}|\textbf{Outcome 4}]$.

If $T_{mut,i}$ is the time to the first mutational event of the $i$th allele (given there is one sensitive cell at $t = 0$) and $T_{og,i}$ is the time to outgrowth for the newly spawned clone of the $i$th allele, then

$$T_{det,i} = T_{mut,i} + T_{og,i} \tag{13}$$

For Alleles A and B, we might expect a timeline that resembles:

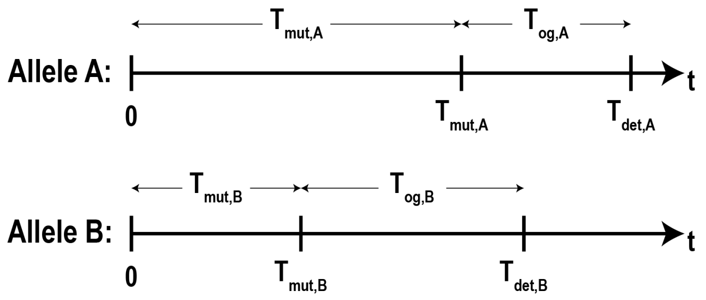


We expect Allele A to grow out faster (i.e. $T_{og,A} < T_{og,B}$) because it is more resistant and thus has a faster growth rate in the presence of drug. However, this does not necessarily mean that it will reach clinical detection first. We see that if an Allele B clone spawns early enough, it will be the dominant allele upon relapse. This is true if $T_{og,B} - T_{og,A} < T_{mut,A} - T_{mut,B}$. Thus,

$$P[\textbf{Outcome 4.B}] = P[T_{det,B} < T_{det,A}] = P[\Delta T_{og} < T_{mut,A} - T_{mut_B}] \tag{14}$$

where $\Delta T_{og} = T_{og,B} - T_{og,A}$.

Because we consider the variation in $T_{og,i}$ to be very small compared to $T_{mut,i}$, we treat $\Delta T_{og}$ as a deterministic variable. Its value can be calculated using resistance allele-specific variations of equation (6):

$$\Delta T_{og} = T_{og,B} - T_{og,A} = \frac{1}{r'_B} \ln\left(\frac{Mr'_B}{b_B}\right) - \frac{1}{r'_A} \ln\left(\frac{Mr'_A}{b_A}\right) \tag{15}$$

From here on, we abbreviate times to mutation $T_{mut,A}$ and $T_{mut,B}$ to $T_A$ and $T_B$, respectively. To obtain a solution for the right-hand expression in equation (14), we find the probability that the difference that these two random variables exceeds a threshold value, $\Delta T_{og}$. The probability that two random variables will take on values on a two-dimensional interval is the integral of their joint probability density function over that interval.

$$P[\Delta T_{og} < T_A - T_B] = \iint_D f^*_{A,B}(t_A, t_B) dA \tag{16}$$

Here, $f^*_{A,B}(t_A, t_B)$ is the conditional joint probability density function for $T_A$ and $T_B$, and $D$ is the region in the first quadrant (where $f^*_{A,B}$ takes on nonzero values) bounded above by the line $t_B = t_A - \Delta T_{og}$. We consider the *conditional* joint probability density function because we are concerned with **Outcome 4**, where both alleles are present.

Because $T_A$ and $T_B$ are independent random variables, their joint conditional probability density function is given by the product of their individual PDFs:

$$f^*_{A,B}(t_A, t_B) = f^*_A(t_A) \cdot f^*_B(t_B) \tag{17}$$

The probability density function for $T_i$, $f_i(t_i)$, gives the probability that a mutational event occurs on the interval $(t_i, t_i + dt_i)$. It is given by the derivative of the cumulative density function $f_i(t_i) = \frac{d}{dt} F_i(t_i)$. Therefore,

$$f_i(t_i) = \frac{d}{dt} F_i(t_i) = \begin{cases} \eta_i \alpha_i e^{r_S t_i} \exp[\frac{-\alpha_i}{r_S} e^{r_S t_i}] & 0 \le t_i < T \\ \nu_i \beta_i e^{r'_S t_i} \exp[\frac{-\beta_i}{r'_S} e^{r'_S t_i}] & T \le t_i \end{cases} \tag{18}$$

To determine the *conditional* probability density function $f^*_i(t_i)$, we divide $f_i(t_i)$ by the probability that the $i$th allele *will* spawn. From equation (10), we have:

$$f^*_i(t_i) = \frac{f_i(t_i)}{F_i(\infty)} = \begin{cases} \frac{\eta_i \alpha_i}{1-\nu_i} e^{r_S t_i} \exp[\frac{-\alpha_i}{r_S} e^{r_S t_i}] & 0 \le t_i < T \\ \frac{\nu_i \beta_i}{1-\nu_i} e^{r'_S t_i} \exp[\frac{-\beta_i}{r'_S} e^{r'_S t_i}] & T \le t_i \end{cases} \tag{19}$$

which is the conditional probability density function of $T_i$ given that at least one mutational event to the $i$th allele eventually takes place. An example plot of $f^*_i(t_i)$ is shown in **Appendix Figure 1.2**.

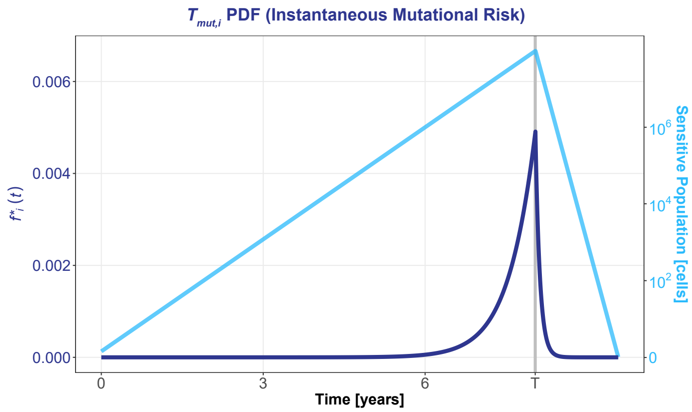


**Appendix Figure 1.2**: Instantaneous mutational risk, conditioned on the mutational event eventually occurring. The same parameters were used as in **Appendix Figure 1.1**.

From equations (17) and (19), we arrive at the formal conditional joint probability density function:

$$f^*_{A,B}(t_A, t_B) = \begin{cases} C_1 \cdot e^{r_S t_A} \exp(\frac{-\alpha_A}{r_S} e^{r_S t_A}) \cdot e^{r_S t_B} \exp(\frac{-\alpha_B}{r_S} e^{r_S t_B}) & 0 \leq t_A, t_B < T \\ C_2 \cdot e^{r_S t_A} \exp(\frac{-\alpha_A}{r_S} e^{r_S t_A}) \cdot e^{r'_S t_B} \exp(\frac{-\beta_B}{r'_S} e^{r'_S t_B}) & 0 \leq t_A < T \leq t_B \\ C_3 \cdot e^{r'_S t_A} \exp(\frac{-\beta_A}{r'_S} e^{r'_S t_A}) \cdot e^{r_S t_B} \exp(\frac{-\alpha_B}{r_S} e^{r_S t_B}) & 0 \leq t_B < T \leq t_A \\ C_4 \cdot e^{r'_S t_A} \exp(\frac{-\beta_A}{r'_S} e^{r'_S t_A}) \cdot e^{r'_S t_B} \exp(\frac{-\beta_B}{r'_S} e^{r'_S t_B}) & T \leq t_A, t_B \end{cases} \quad (20)$$

The simplifying constants $C_j$ are defined as follows:

$$\begin{aligned} C_1 &= \frac{\alpha_A \eta_A}{1-\nu_A} \frac{\alpha_B \eta_B}{1-\nu_B} \\ C_2 &= \frac{\alpha_A \eta_A}{1-\nu_A} \frac{\beta_B \nu_B}{1-\nu_B} \\ C_3 &= \frac{\beta_A \nu_A}{1-\nu_A} \frac{\alpha_B \eta_B}{1-\nu_B} \\ C_4 &= \frac{\beta_A \nu_A}{1-\nu_A} \frac{\beta_B \nu_B}{1-\nu_B} \end{aligned} \quad (21)$$

An example joint conditional PDF $f^*_{A,B}(t_A, t_B)$ is shown in **Appendix Figure 1.3**.

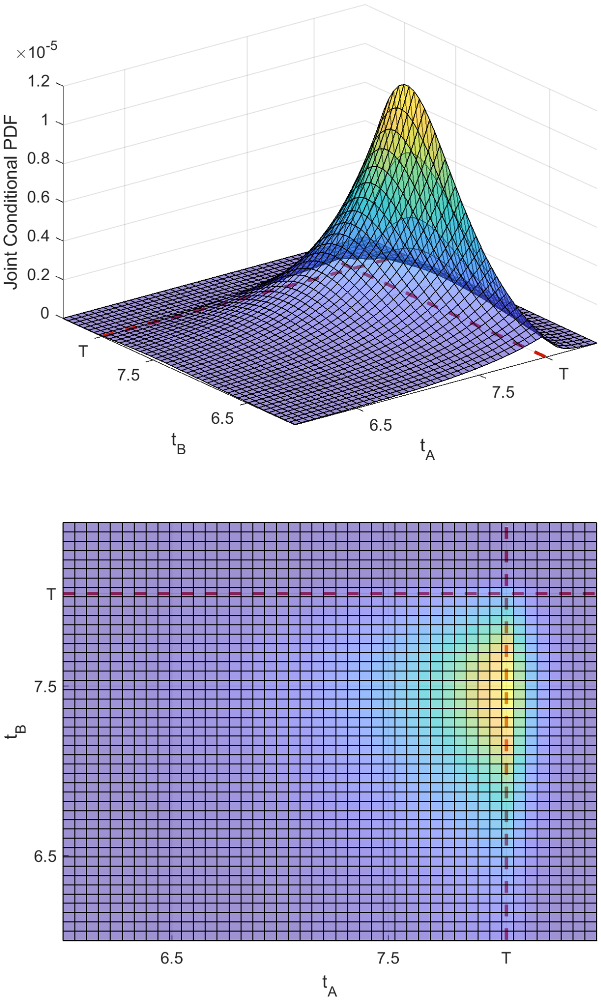


**Appendix Figure 1.3**: Joint conditional PDF of $T_A$ and $T_B$. For $T_A$, the same parameters were used as in **Appendix Figure 1.3** (E255V). For $T_B$, $\rho_B = 0.97$ and $r'_B = 0.0042$ to match parameters for E255K.

Note that $f^*_{A,B}(t_A, t_B)$ is a piecewise function with four subdomains. Thus, when evaluating the integral in equation (16), it is necessary to separate it into appropriate subintegrals. The choice of subintegral and their associated limits of integration depend on the value of $\Delta T_{og}$. From here on, we abbreviate $\Delta T_{og}$ to $\Delta T$. Given our assumption that Allele A is more resistant than Allele B, $\Delta T = T_{og,B} - T_{og,A}$ must be positive. Consider the two cases:

**Case 1**: $0 \leq \Delta T < T$

**Case 2**: $T \leq \Delta T$

We begin with **Case 1**.

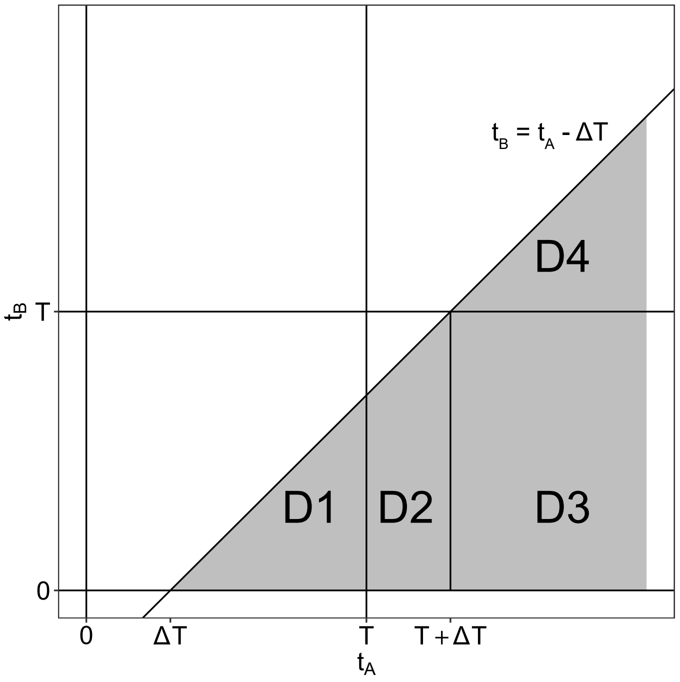


**Appendix Figure 1.4**: $t_A$-$t_B$ plane for **Case 1**: $0 \leq \Delta T < T$. The shaded area represents the region where the condition $\Delta T < T_A - T_B$ (equation (16)) is met.

The region of integration is divided into four subregions in the $t_A$-$t_B$ plane:

$$\iint_D f^* dA = \iint_{D_1} f^* dA + \iint_{D_2} f^* dA + \iint_{D_3} f^* dA + \iint_{D_4} f^* dA \qquad (22)$$

We now take the integral for each individual subregion. For $D_1$ we have:

$$\begin{aligned}
\iint_{D_1} f^* dA &= C_1 \int_{\Delta T}^{T} e^{r_S t_A} \exp(-\frac{\alpha_A}{r_S} e^{r_S t_A}) \int_0^{t_A - \Delta T} e^{r_S t_B} \exp(-\frac{\alpha_B}{r_S} e^{r_S t_B}) dt_B dt_A \\
&= -\frac{C_1}{\alpha_B} \int_{\Delta T}^{T} e^{r_S t_A} \exp(-\frac{\alpha_A}{r_S} e^{r_S t_A}) \Big[ \exp(-\frac{\alpha_B}{r_S} e^{-r_S \Delta T} e^{r_S t_A)}) - \exp(-\frac{\alpha_B}{r_S}) \Big] dt_A \\
&= -\frac{C_1}{\alpha_B} \int_{\Delta T}^{T} e^{r_S t_A} \exp\Big( (-\frac{1}{r_S}(\alpha_A + \alpha_B e^{-r_S \Delta T}) e^{r_S t_A} \Big) dt_A + \\
&\frac{C_1}{\alpha_B} e^{-\alpha_B / r_S} \int_{\Delta T}^{T} e^{r_S t_A} \exp(-\frac{\alpha_A}{r_S} e^{r_S t_A}) dt_A \\
&= \frac{C_1}{\alpha_B(\alpha_A + \alpha_B e^{-r_S \Delta T})} \Big[ \exp\Big( -\frac{e^{r_S T}}{r_S}(\alpha_A + \alpha_B e^{-r_S \Delta T}) \Big) - \exp\Big( -\frac{e^{r_S \Delta T}}{r_S}(\alpha_A + \alpha_B e^{-r_S \Delta T}) \Big) \Big] - \\
&\frac{C_1}{\alpha_A \alpha_B} e^{-\alpha_B / r_S} \Big[ \exp(-\frac{\alpha_A}{r_S} e^{r_S T}) - \exp(-\frac{\alpha_A}{r_S} e^{r_S \Delta T}) \Big]
\end{aligned} \tag{23}$$

For $D_2$:

$$\begin{aligned}
\iint_{D_2} f^* dA &= C_3 \int_{T}^{T+\Delta T} e^{r'_S t_A} \exp(-\frac{\beta_A}{r'_S} e^{r'_S t_A}) \int_0^{t_A - \Delta T} e^{r_S t_B} \exp(-\frac{\alpha_B}{r_S} e^{r_S t_B}) dt_B dt_A \\
&= -\frac{C_3}{\alpha_B} \int_{T}^{T+\Delta T} e^{r'_S t_A} \exp(-\frac{\beta_A}{r'_S} e^{r'_S t_A}) \cdot \Big[ \exp(-\frac{\alpha_B}{r_S} e^{-r_S \Delta T} e^{r_S t_A)}) - \exp(-\frac{\alpha_B}{r_S}) \Big] dt_A \\
&= -\frac{C_3}{\alpha_B} \int_{T}^{T+\Delta T} e^{r'_S t_A} \exp(-\frac{\beta_A}{r'_S} e^{r'_S t_A}) \exp(-\frac{\alpha_B}{r_S} e^{-r_S \Delta T} e^{r_S t_A}) dt_A + \\
&\frac{C_3}{\alpha_B} e^{-\alpha_B / r_S} \int_{T}^{T+\Delta T} e^{r'_S t_A} \exp(-\frac{\beta_A}{r'_S} e^{r'_S t_A}) dt_A
\end{aligned} \tag{24}$$

The second integral in the last form of equation (24) has a straightforward solution:

$$\begin{aligned}
&\frac{C_3}{\alpha_B} e^{-\alpha_B / r_S} \int_{T}^{T+\Delta T} e^{r'_S t_A} \exp(-\frac{\beta_A}{r'_S} e^{r'_S t_A}) dt_A = \\
&\qquad -\frac{C_3}{\beta_A \alpha_B} e^{-\alpha_B / r_S} \Big[ \exp(-\frac{\beta_A}{r'_S} e^{r'_S (T+\Delta T)}) - \exp(-\frac{\beta_A}{r'_S} e^{r'_S T}) \Big]
\end{aligned} \tag{25}$$

However, the first integral in the last form of equation (24) requires more analysis. Let $u = e^{r'_S t_A}$, $du = r'_S e^{r'_S t_A} dt_A$, $u_1 = e^{r'_S T}$, and $u_2 = e^{r'_S (T+\Delta T)}$.

Now, using the power series for exponential functions:

$$
\begin{aligned}
-&\frac{C_3}{\alpha_B}\int_T^{T+\Delta T} e^{r'_S t_A}\exp(-\frac{\beta_A}{r'_S}e^{r'_S t_A})\exp(-\frac{\alpha_B}{r_S}e^{-r_S\Delta T}e^{r_S t_A})dt_A \\
&= -\frac{C_3}{\alpha_B r'_S}\int_{u_1}^{u_2}\exp(-\frac{\beta_A}{r'_S}u)\exp(-\frac{\alpha_B}{r_S}e^{-r_S\Delta T}u^{r_S/r'_S})du \\
&= -\frac{C_3}{\alpha_B r'_S}\int_{u_1}^{u_2}\sum_{k=0}^{\infty}\frac{(-u\beta_A/r'_S)^k}{k!}\exp(-\frac{\alpha_B}{r_S}e^{-r_S\Delta T}u^{r_S/r'_S})du \\
&= -\frac{C_3}{\alpha_B r'_S}\sum_{k=0}^{\infty}\frac{(-\beta_A/r'_S)^k}{k!}\int_{u_1}^{u_2}\exp(-\frac{\alpha_B}{r_S}e^{-r_S\Delta T}u^{r_S/r'_S})u^k du
\end{aligned}
\tag{26}
$$

Let $v = \frac{\alpha_B}{r_S}e^{-r_S\Delta T}u^{r_S/r'_S}$, $dv = \frac{\alpha_B}{r'_S}e^{-r_S\Delta T}u^{(r_S/r'_S-1)}du$, $v_1 = \frac{\alpha_B}{r_S}e^{r_S(T-\Delta T)}$ and $v_2 = \frac{\alpha_B}{r_S}e^{r_S T}$. Solving for $u$ and $du$, we have $u = (\frac{r_S}{\alpha_B}e^{r_S\Delta T}v)^{r'_S/r_S}$ and $du = \frac{r'_S}{r_S}(\frac{r_S}{\alpha_B}e^{r_S\Delta T})^{r'_S/r_S}v^{(r'_S/r_S-1)}dv$. Now,

$$
\begin{aligned}
-&\frac{C_3}{\alpha_B}\int_T^{T+\Delta T} e^{r'_S t_A}\exp(-\frac{\beta_A}{r'_S}e^{r'_S t_A})\exp(-\frac{\alpha_B}{r_S}e^{-r_S\Delta T}e^{r_S t_A})dt_A \\
&= -\frac{C_3}{\alpha_B r'_S}\sum_{k=0}^{\infty}\frac{(-\beta_A/r'_S)^k}{k!}\int_{v_1}^{v_2} e^{-v}(\frac{r_S}{\alpha_B}e^{r_S\Delta T}v)^{kr'_S/r_S}\cdot\frac{r'_S}{r_S}(\frac{r_S}{\alpha_B}e^{r_S\Delta T})^{r'_S/r_S}v^{r'_S/r_S-1}dv \\
&= -\frac{C_3}{\alpha_B r_S}(\frac{r_S}{\alpha_B}e^{r_S\Delta T})^{r'_S/r_S}\sum_{k=0}^{\infty}\frac{(-\beta_A/r'_S)^k}{k!}(\frac{r_S}{\alpha_B}e^{r_S\Delta T})^{kr'_S/r_S}\int_{v_1}^{v_2} e^{-v}v^{(kr'_S/r_S+r'_S/r_S-1)}dv
\end{aligned}
\tag{27}
$$

Finally, using the integral definition of the lower incomplete gamma function $\gamma(a,x) = \int_0^x t^{a-1}e^{-t}dt$, we arrive at an asymptotic solution for this integral:

$$
\begin{aligned}
-&\frac{C_3}{\alpha_B}\int_T^{T+\Delta T} e^{r'_S t_A}\exp(-\frac{\beta_A}{r'_S}e^{r'_S t_A})\exp(-\frac{\alpha_B}{r_S}e^{-r_S\Delta T}e^{r_S t_A})dt_A \\
&= K\sum_{k=0}^{\infty}\Bigg\{\frac{(-\beta_A/r'_S)^k}{k!}(\frac{r_S}{\alpha_B}e^{r_S\Delta T})^{kr'_S/r_S}\cdot \\
&\qquad \Big[\gamma(\frac{r'_S}{r_S}(k+1),\frac{\alpha_B}{r_S}e^{r_S T}) - \gamma(\frac{r'_S}{r_S}(k+1),\frac{\alpha_B}{r_S}e^{r_S(T-\Delta T)})\Big]\Bigg\}
\end{aligned}
\tag{28}
$$

where $K = -\frac{C_3}{\alpha_B r_S}(\frac{r_S}{\alpha_B}e^{r_S\Delta T})^{r'_S/r_S}$. Given that the error bound on the series in equation (28) is the $(k+1)$th term of the series, we can evaluate the integral to a desired degree of precision.

For $D_3$:

$$
\begin{aligned}
\iint_{D_3} f^* dA = C_3 \int_{T+\Delta T}^{\infty} e^{r'_S t_A} \exp(-\frac{\beta_A}{r'_S} e^{r'_S t_A}) dt_A \int_0^T e^{r_S t_B} \exp(-\frac{\alpha_B}{r_S} e^{r_S t_B}) dt_B \\
= \frac{C_3}{\beta_A \alpha_B} \Big[1 - \exp(-\frac{\beta_A}{r'_S} e^{r'_S (T+\Delta T)})\Big] \cdot \Big[\exp(-\frac{\alpha_B}{r_S} e^{r_S T}) - \exp(-\frac{\alpha_B}{r_S})\Big]
\end{aligned}
\tag{29}
$$

For $D_4$:

$$
\begin{aligned}
\iint_{D_4} f^* dA &= C_4 \int_{T+\Delta T}^{\infty} e^{r'_S t_A} \exp(-\frac{\beta_A}{r'_S} e^{r'_S t_A}) \int_T^{t_A - \Delta T} e^{r'_S t_B} \exp(-\frac{\beta_B}{r'_S} e^{r'_S t_B}) dt_B dt_A \\
&= -\frac{C_4}{\beta_B} \int_{T+\Delta T}^{\infty} e^{r'_S t_A} \exp(-\frac{\beta_A}{r'_S} e^{r'_S t_A}) [\exp(-\frac{\beta_B}{r'_S} e^{-r'_S \Delta T} e^{r'_S t_A}) - \exp(-\frac{\beta_B}{r'_S} e^{r'_S T})] dt_A \\
&= -\frac{C_4}{\beta_B} \int_{T+\Delta T}^{\infty} e^{r'_S t_A} \exp\Big(-\frac{e^{r'_S t_A}}{r'_S} (\beta_A + \beta_B e^{-r'_S \Delta T})\Big) dt_A + \\
&\frac{C_4}{\beta_B} \exp(-\frac{\beta_B}{r'_S} e^{r'_S T}) \int_{T+\Delta T}^{\infty} e^{r'_S t_A} \exp(-\frac{\beta_A}{r'_S} e^{r'_S t_A}) dt_A \\
&= \frac{C_4}{\beta_B (\beta_A + \beta_B e^{-r'_S \Delta T})} \Big[1 - \exp\Big(-\frac{e^{r'_S (T+\Delta T)}}{r'_S} (\beta_A - \beta_B e^{-r'_S \Delta T})\Big)\Big] - \\
&\frac{C_4}{\beta_A \beta_B} \exp(-\frac{\beta_B}{r'_S} e^{r'_S T}) \Big[1 - \exp(-\frac{\beta_A}{r'_S} e^{r'_S (T+\Delta T)})\Big]
\end{aligned}
\tag{30}
$$

Thus, the full expression for equation (22) is:

$$\iint_D f^* dA = \frac{C_1}{\alpha_B(\alpha_A + \alpha_B e^{-r_S \Delta T})} \Big[ \exp\Big( -\frac{e^{r_S T}}{r_S}(\alpha_A + \alpha_B e^{-r_S \Delta T})\Big) -$$
$$\exp\Big( -\frac{e^{r_S \Delta T}}{r_S}(\alpha_A + \alpha_B e^{-r_S \Delta T})\Big)\Big] -$$
$$\frac{C_1}{\alpha_A \alpha_B} e^{-\alpha_B/r_S} \Big[ \exp(-\frac{\alpha_A}{r_S} e^{r_S T}) - \exp(-\frac{\alpha_A}{r_S} e^{r_S \Delta T})\Big] +$$
$$K \sum_{k=0}^{\infty} \Bigg\{ \frac{(-\beta_A/r'_S)^k}{k!} (\frac{r_S}{\alpha_B} e^{r_S \Delta T})^{k r'_S / r_S} \cdot$$
$$\Big[ \gamma(\frac{r'_S}{r_S}(k+1), \frac{\alpha_B}{r_S} e^{r_S T}) - \gamma(\frac{r'_S}{r_S}(k+1), \frac{\alpha_B}{r_S} e^{r_S (T - \Delta T)})\Big] \Bigg\} - \tag{31}$$
$$\frac{C_3}{\beta_A \alpha_B} e^{-\alpha_B/r_S} \Big[ \exp(-\frac{\beta_A}{r'_S} e^{r'_S (T+\Delta T)}) - \exp(-\frac{\beta_A}{r'_S} e^{r'_S T})\Big] +$$
$$\frac{C_3}{\beta_A \alpha_B} \Big[ 1 - \exp(-\frac{\beta_A}{r'_S} e^{r'_S (T+\Delta T)})\Big] \cdot \Big[ \exp(-\frac{\alpha_B}{r_S} e^{r_S T}) - \exp(-\frac{\alpha_B}{r_S})\Big] +$$
$$\frac{C_4}{\beta_B(\beta_A + \beta_B e^{-r_S \Delta T})} \Big[ 1 - \exp\Big( -\frac{e^{r'_S (T+\Delta T)}}{r'_S}(\beta_A - \beta_B e^{-r'_S \Delta T})\Big)\Big] -$$
$$\frac{C_4}{\beta_A \beta_B} \exp(-\frac{\beta_B}{r'_S} e^{r'_S T}) \Big[ 1 - \exp(-\frac{\beta_A}{r_S} e^{r'_S (T+\Delta T)})\Big]$$

Similarly, it can be shown that $\iint_D f^* dA$ for **Case 2** ($T \le \Delta T$) is:

$$\iint_D f^* dA = K \sum_{k=0}^{\infty} \Bigg\{ \frac{(-\beta_A/r'_S)^k}{k!} (\frac{r_S}{\alpha_B} e^{r_S \Delta T})^{k r'_S / r_S} \cdot$$
$$\Big[ \gamma(\frac{r'_S}{r_S}(k+1), \frac{\alpha_B}{r_S} e^{r_S T}) - \gamma(\frac{r'_S}{r_S}(k+1), \frac{\alpha_B}{r_S})\Big] \Bigg\} -$$
$$\frac{C_3}{\beta_A \alpha_B} e^{-\alpha_B/r_S} \Big[ \exp\Big( -\frac{\beta_A}{r'_S} e^{r'_S (T+\Delta T)}\Big) - \exp\Big( -\frac{\beta_A}{r'_S} e^{r'_S \Delta T}\Big)\Big] +$$
$$\frac{C_3}{\beta_A \alpha_B} \Big[ 1 - \exp\Big( -\frac{\beta_A}{r'_S} e^{r'_S (T+\Delta T)}\Big)\Big] \cdot \Big[ \exp\Big( -\frac{\alpha_B}{r_S} e^{r_S T}\Big) - \exp\Big( -\frac{\alpha_B}{r_S}\Big)\Big] +$$
$$\frac{C_4}{\beta_B(\beta_A + \beta_B e^{-r'_S \Delta T})} \Big[ 1 - \exp\Big( -\frac{e^{r'_S (T+\Delta T)}}{r'_S}(\beta_A + \beta_B e^{-r'_S \Delta T})\Big)\Big] -$$
$$\frac{C_4}{\beta_A \beta_B} \exp(-\frac{\beta_B}{r'_S} e^{r'_S T}) \Big[ 1 - \exp\Big( -\frac{\beta_A}{r'_S} e^{r'_S (T+\Delta T)}\Big)\Big] \tag{32}$$

### 5 Conclusion

In summary, the probability that either allele is dominant upon relapse is the sum of the probability of its fixation and the probability of it outcompeting the other allele (equation (1)).

The probability of each allele's fixation is given by equations (10) and (11):

$$\begin{aligned} P[\text{Allele A is fixed}] &= P[\textbf{Outcome 2}] = (1-\nu_A)\cdot\nu_B \\ P[\text{Allele B is fixed}] &= P[\textbf{Outcome 3}] = \nu_A\cdot(1-\nu_B) \end{aligned} \tag{33}$$

The probability of either allele outcompeting the other allele is given by equations (11), (14), and (16):

$$\begin{aligned} P[\text{Allele A detected first}] &= P[\textbf{Outcome 4}]\cdot P[\textbf{Outcome 4.A}|\textbf{Outcome 4}] \\ &= (1-\nu_A)\cdot(1-\nu_B)\cdot\Big(1-\iint_D f^*_{A,B}(t_A,t_B)dA\Big) \\ P[\text{Allele B detected first}] &= P[\textbf{Outcome 4}]\cdot P[\textbf{Outcome 4.B}|\textbf{Outcome 4}] \\ &= (1-\nu_A)\cdot(1-\nu_B)\cdot\iint_D f^*_{A,B}(t_A,t_B)dA \end{aligned} \tag{34}$$

where $\iint_D f^*_{A,B}(t_A,t_B)dA$ is given by equation (29) for **Case 1** ($0\le\Delta T<T$) and equation (30) for **Case 2** ($T\le\Delta T$).

Thus, the probability that either allele is dominant upon relapse is given by substituting equations (33) and (34) into equation (1):

$$\begin{aligned} &P[\text{Allele A is dominant upon relapse}] \\ &\quad= P[\textbf{Outcome 2}] + P[\textbf{Outcome 4}]\cdot P[\textbf{Outcome 4.A}|\textbf{Outcome 4}] \\ &\quad= (1-\nu_A)\cdot\nu_B + (1-\nu_A)\cdot(1-\nu_B)\cdot\Big(1-\iint_D f^*_{A,B}(t_A,t_B)dA\Big) \\ &P[\text{Allele B is dominant upon relapse}] \\ &\quad= P[\textbf{Outcome 3}] + P[\textbf{Outcome 4}]\cdot P[\textbf{Outcome 4.B}|\textbf{Outcome 4}] \\ &\quad= (1-\nu_B)\cdot\nu_A + (1-\nu_A)\cdot(1-\nu_B)\cdot\iint_D f^*_{A,B}(t_A,t_B)dA\cdot \end{aligned} \tag{35}$$
