## Supplemental Appendix 2 for "Multi-scale predictions of drug resistance epidemiology suggest a new design principle for rational drug design"

### Supplemental Appendix S2:
Stochastic CML Model

Leighow and Liu et al.

July 14, 2019

### 1 Introduction

We developed a stochastic model of evolutionary dynamics in chronic myelogenous leukemia to test the hypothesis that accounting for mutation bias adds predictive value to determining the frequency of specific resistance mutations.

### 2 Pharmacokinetic Model

Towards identifying appropriate drug kill terms for each drug/genotype combination, we used clinical pharmacokinetic data and *in vitro* dose response assays with BaF3 cells to predict the death rate of leukemic cells in response to three BCR-ABL inhibitors: imatinib, nilotinib, and dasatinib.

We begin with the pharmacokinetic simulations. Patient pharmacokinetic profiles are heterogeneous; it is necessary to consider the variation in drug concentration across patients to adequately parameterize the drug kill rates in our model. Furthermore, the relationship between dose and cell killing is nonlinear, and so $C_{ave}$ alone is insufficient to predict drug killing. For this reason, it is necessary to consider temporal fluctuations in drug concentration.

#### 2.1 Pharmacokinetic Parameterization

To generate accurate drug kill terms for our virtual patients, we began by assigning $C_{min}$ and $C_{max}$ values drawn from clinically reported distributions (imatinib [**1**], nilotinib [**2**], and dasatinib [**3**]). To determine the drug concentration as a function of time, we assumed a model of drug absorption and elimination with two compartments: gut and periphery. The ODEs governing the dynamics are:

$$\begin{aligned}\frac{dx}{dt} &= -ax \\ \frac{dy}{dt} &= ax - by\end{aligned} \tag{1}$$

where $x(t)$ and $y(t)$ are the concentrations of drug in the gut and periphery, respectively, $a$ is an absorption rate, and $b$ is an elimination rate. This system of ODEs has a closed form solution for the peripheral drug concentration $y(t)$ for initial conditions $x(0) = x_0$ and $y(0) = y_0$ (using Maple's *dsolve*):

$$y(t) = \left(\frac{ax_0}{a-b} + y_0\right)e^{-bt} - \frac{ax_0}{a-b}e^{-at} \tag{2}$$

To fit the parameters $a$, $b$, $x_0$, and $y_0$ to the generated $C_{min}$ and $C_{max}$ values for each virtual patient and the known $t_{peak}$ for each drug, we used a numerical equation solver (MATLAB's *fsolve*; the trust-region dogleg algorithm) to identify the rate constants and initial conditions that satisfy the definitions of $C_{min}$ and $C_{max}$. At steady state, the following conditions must be met:

$$\begin{aligned} y(0) &= C_{min} \\ y(t_{peak}) &= C_{max} \\ y'(t_{peak}) &= 0 \\ y(t_{dose}) &= C_{min} \end{aligned} \tag{3}$$

By solving for pharmacokinetic parameters $a$, $b$, $x_0$, and $y_0$ that satisfy this system of equations, we arrive at a function $y(t)$ which describes the peripheral drug concentration as a function of time. An example pharmacokinetic profile is shown in **Appendix Figure 2.1**.

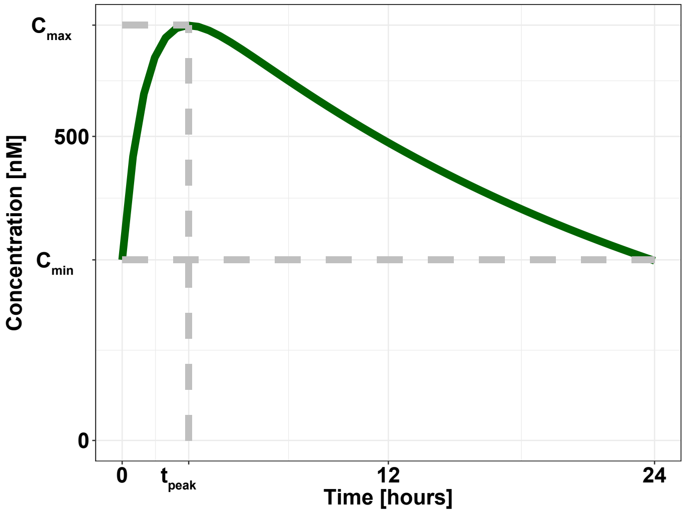


**Appendix Figure 2.1**: Example trajectory of imatinib pharmacokinetics. Here, $C_{max} = 683$ nM, $C_{min} = 297$ nM, and $t_{peak} = 3$ hours. The corresponding pharmacokinetic parameters are $a = 22.1$ /hr, $b = 1.01$ /hr, $x_0 = 491$ nM, and $y_0 = 297$ nM.

For each drug, $10^4$ virtual patients were simulated.

#### 2.2 Drug Kill Rate

Next, we determined the drug kill term as a function of concentration. Using our drug dose response values for BaF3s, we fit a two parameter Hill equation for each drug/genotype combination:

$$z = \frac{1}{1 + \Big(y/C\Big)^h} \tag{4}$$

where $z$ is the fraction of cells surviving, $y$ is the drug concentration, and the parameters $C$ and $h$ are the $IC_{50}$ and Hill coefficient, respectively.

To determine the rate of drug killing $\alpha$, we used the formula previously derived by Zhao et al (2016) [**4**]:

$$\alpha = \frac{-\ln(z)}{t_{assay}} \tag{5}$$

Here, $t_{assay}$ is 72 hours (see Methods).

By substituting the function $y(t)$ into equations (4) and (5), we arrive at $\alpha(t)$, the drug kill term as a function of time given our pharmacokinetic profile.

$$\alpha(t) = \frac{-1}{t_{assay}} \ln\left(\frac{1}{1 + (\Big(y(t)/C\Big)^h}\right) \tag{6}$$

We take $\alpha_{ave}$, the mean value of $\alpha(t)$ over the dosing period, to be the effective drug kill rate for each drug/genotype combination in the BaF3 system.

#### 2.3 Parameterizing Leukemic Stem Cell System

Since our *in vitro* and *in vivo* systems have different dynamics, we transform the drug kill rate from the *in vitro* BaF3 system to the leukemic stem cell system used in our stochastic model described below.

For drug kill rates measured in BaF3 cells expressing wild-type BCR-ABL, we linearly transformed the $\alpha_{ave}$ values such that the mean value agreed with the mean drug kill rate reported by Fassoni et al (2018) [**5**].

Given the low incidence of CML, the rates of clinical outgrowth for each resistant variant have not been measured. Therefore, we linearly transformed the $\alpha_i$ values assuming equivalent ratios for net growth rates ($g$) and drug kill rates ($\alpha$) to preserve the qualitative phenotype of resistance:

$$\frac{g_{LSC}}{g_{BaF3}} = \frac{\alpha_{LSC,i}}{\alpha_{BaF3,i}} \tag{7}$$

For our analysis, we removed simulated patients who were not predicted to respond to therapy (i.e. cases where the drug kill rate for WT cells did not exceed the net growth rate), since genetic resistance has no clinical significance for a patient that does not respond to treatment.

### 3 Stochastic Dynamic Model

We simulated the evolutionary trajectory of each virtual patient's leukemia as a continuous-time birth-death process. The model was implemented using MATLAB (GitHub: `https://github.com/pritchardlabatpsu/SurvivalOfTheLikeliest/tree/master/Figures/Figure4`). The stochastic solution was approximated using a Gillespie algorithm modified with adaptive tau leaping (Cao et al (2006) [**6**]) to improve computational efficiency.

#### 3.1 Leukemic Stem Cell Model

We adapted a model of leukemic stem cell (LSC) dynamics from Fassoni et al (2018) [**5**] to include resistance mutations. The original model was parameterized using phase 3 clinical trial data (N = 122). The basic model includes three compartments: quiescent LSCs (QLSC: $X$), proliferating LSCs (PLSC: $Y$), and differentiated leukemic cells (DLC: $Z$). Quiescent LSCs are activated at a rate of $p_{XY}$. Proliferating LSCs are deactivated at a rate of $p_{YX}$, divide at a net growth rate of $p_Y$, and differentiate at a rate of $p_W$. The proliferating LSC net growth rate $p_Y$ was deconstructed into division and death rates, $b_Y$ and $d_Y$, respectively. Assuming a turnover rate $r_{to} = 0.3$ (estimate from Komarova and Wodarz (2005) [**7**]), the rates can be related according to the following equations:

$$\begin{aligned} b_Y &= \frac{p_Y}{1 - r_{to}} \\ d_Y &= b_Y - p_Y \end{aligned} \tag{8}$$

DLCs die at a rate of $r_W$. The model assumes that the TKI targets proliferating LSCs (an assumption supported by the biphasic molecular response observed in the clinic, as described previously [**8**],[**9**]) at a variant-specific rate of $\alpha_i$. Resistant proliferating LSCs are generated from the sensitive wild-type proliferating LSCs at a rate of $\rho_i \mu$, where $\rho_i$ is a variant-specific conditional mutational probability (given a resistance mutation occurs) and $\mu$ is the resistance mutation rate.

The model is summarized by the propensity function $a(\mathbf{x})$ and state change matrix $\nu$:

$$a(\mathbf{x}) = \frac{1}{a_0} \begin{bmatrix} p_{XY}X_{WT} \\ p_{YX}Y_{WT} \\ b_Y(1-\mu)Y_{WT} \\ (d_Y+\alpha_{WT})Y_{WT} \\ p_W Y_{WT} \\ r_W W_{WT} \\ \rho_{M244V}\mu b_Y Y_{WT} \\ p_{XY}X_{M244V} \\ p_{YX}Y_{M244V} \\ b_Y Y_{M244V} \\ (d_Y+\alpha_{M244V})Y_{M244V} \\ p_W Y_{M244V} \\ r_W W_{M244V} \\ \vdots \end{bmatrix} \tag{9}$$

$$\nu = \begin{bmatrix} -1 & 1 & 0 & 0 & 0 & 0 & \dots \\ 1 & 1 & 0 & 0 & 0 & 0 & \dots \\ 0 & 1 & 0 & 0 & 0 & 0 & \dots \\ 0 & -1 & 0 & 0 & 0 & 0 & \dots \\ 0 & 0 & 1 & 0 & 0 & 0 & \dots \\ 0 & 0 & 0 & 0 & 0 & -1 & \dots \\ 0 & 0 & 0 & 0 & 1 & 0 & \dots \\ 0 & 0 & 0 & -1 & 1 & 0 & \dots \\ 0 & 0 & 0 & 1 & -1 & 0 & \dots \\ 0 & 0 & 0 & 0 & 1 & 0 & \dots \\ 0 & 0 & 0 & 0 & -1 & 0 & \dots \\ 0 & 0 & 0 & 0 & 0 & 1 & \dots \\ 0 & 0 & 0 & 0 & 0 & -1 & \dots \\ \vdots & \vdots & \vdots & \vdots & \vdots & \vdots & \ddots \end{bmatrix} \tag{10}$$

Here, $a_0$ is the sum of the rates in the propensity function.

##### 3.2 Simulation Conditions

At $t = 0$ there exist only 50 wild-type proliferating LSCs ($Y_{WT,0} = 50$). Initially, $\alpha = 0$ for all variants. Once the differentiated leukemic cell population reaches detection, $\alpha$ values were set to the appropriate allele-specific drug kill term as described above. In the clinic, the size of leukemic population at detection is variable; thus, each simulated patient was assigned a detection size drawn from a clinically-parameterized lognormal distribution (Stein et al (2011) [**8**]). Simulations continued until either a resistant variant spawned and reached detection or the wild-type proliferating LSC population was eradicated.

For each drug, simulations were run twice - once with mutation bias and once without. With mutation bias, $\rho_i$ was an allele-specific parameter representing the relative probability of a mutation to that specific variant (see Methods).

Without mutation bias, $\rho_i$ was identical for each variant (i.e. $\rho_i = \frac{1}{n}$ where $n$ is the number of resistance variants). The resistance mutation rate $\mu$ was set to $4 \times 10^{-8}$ as in Michor et al (2005) [**9**].

#### 3.3 Model Parameters

Model parameters are outlined in the tables below.

**Appendix Table 2.1**: General Model Parameters

| **Parameter** | **Description** | **Value** | **Unit** |
|---|---|---|---|
| $p_{XY}$ | QLSC activation rate | 0.0018 | [/day] |
| $p_{YX}$ | PLSC deactivation rate | $3.13 \times 10^{-5}$ | [/day] |
| $b_Y$ | PLSC division rate | 0.0088 | [/day] |
| $d_Y$ | PLSC death rate | 0.0026 | [/day] |
| $\alpha_{WT}$ | Wild-type PLSC drug kill rate | See Table () | [/day] |
| $p_W$ | PLSC differentiation rate | $1.32 \times 10^5$ | [/day] |
| $r_W$ | DLC death rate | 0.13 | [/day] |
| $\mu$ | Resistance mutation rate | $4 \times 10^{-8}$ | [/division] |
| $\rho_i$ | Allele-specific mutation probability | See **Table 2.2** | [/mutation] |
| $\alpha_i$ | Allele-specific drug kill rate | See **Table 2.2** | [/day] |

Mutation probabilities $\rho_i$ and mean drug kill rates $\alpha_i$ are given in the table below.

**Appendix Table 2.2**: Allele-Specific Parameters

| **Variant** | $\rho_\mathbf{i}$ | **Imatinib** $\alpha_\mathbf{i}$ | **Nilotinib** $\alpha_\mathbf{i}$ | **Dasatinib** $\alpha_\mathbf{i}$ |
|---|---|---|---|---|
| WT | – | 0.049 | 0.18 | 0.19 |
| M244V | 0.038 | 0.0063 | 0.011 | 0.0080 |
| L248V | 0.012 | 0.00093 | 0.0041 | 0.0035 |
| G250E | 0.13 | 0.00028 | 0.0022 | 0.0040 |
| Q252H | 0.023 | 0.00051 | 0.0056 | 0.0064 |
| Y253F | 0.0046 | 0.00072 | 0.0056 | 0.013 |
| Y253H | 0.055 | $5.10\times10^{-5}$ | $6.40\times10^{-5}$ | 0.0068 |
| E255K | 0.13 | 0.00045 | 0.0034 | 0.0068 |
| E255V | 0.0046 | $7.7\times10^{-10}$ | 0.00011 | 0.0040 |
| D276G | 0.038 | 0.0047 | 0.0091 | 0.012 |
| T315I | 0.18 | $1.4\times10^{-10}$ | $1.410^{-11}$ | $1.2\times10^{-11}$ |
| F317L | 0.077 | 0.0027 | 0.0054 | 0.0032 |
| M351T | 0.055 | 0.0020 | 0.010 | 0.013 |
| E355G | 0.038 | 0.0025 | 0.0087 | 0.009 |
| F359C | 0.0092 | 0.00066 | 0.0025 | 0.011 |
| F359I | 0.0031 | 0.0029 | 0.0027 | 0.0081 |
| F359V | 0.0092 | 0.0022 | 0.0029 | 0.0073 |
| H396P | 0.011 | 0.00024 | 0.0023 | 0.0071 |
| H396R | 0.038 | 0.0025 | 0.0028 | 0.0021 |
| E459K | 0.13 | 0.0047 | 0.0094 | 0.0090 |
