## Supplemental Appendix 3 for "Multi-scale predictions of drug resistance epidemiology suggest a new design principle for rational drug design"

### Supplemental Appendix S3:
#### Adjuvant Therapy Model

Leighow and Liu et al.

July 14, 2019

##### 0.1 Introduction

Adjuvant therapy is the practice of treating a tumor in conjunction with surgery. The expectation is that surgery removes the bulk of the tumor, while the adjuvant therapy removes cancer cells that escaped resection. We simulated a cohort of virtual patients that began with a small population of drug-sensitive cancer cells. Upon detection of the disease, a large portion of the tumor is removed and the remaining sensitive cells are treated with drug (such that their net growth rate is negative) until the sensitive population is eradicated. Over the lifespan of the cancer, sensitive cells have the potential to seed resistant subclones (10 possible variants simulated with allele-specific resistances and biases) with positive net growth rates in the presence of drug. The simulation continues until eradication of all cells or progression due to drug resistance, at which point the dominant resistance allele is noted.

##### 0.2 Parameters

For all genotypes, division rates and natural death rates were $b = 0.14$ and $d = 0.13$ /day, respectively. The drug kill rate for sensitive cells was $\alpha_S = 0.04$ /day [**1**]. Allele-specific drug kill rates for resistance variants were drawn from an exponential distribution to reflect the distribution of resistance in CML (**Appendix Figure 3.1**).

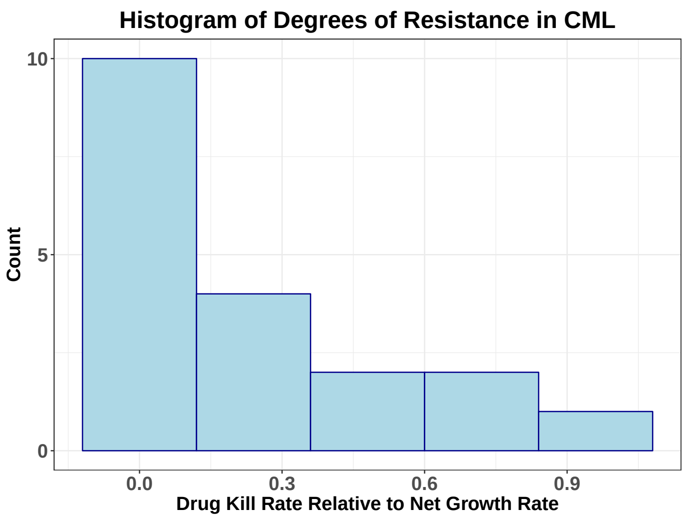


**Appendix Figure 3.1**: Histogram of degrees of resistance in CML. Values on the x-axis are drug kill rates for imatinib-resistant BCR-ABL variants in BaF3 cells (as calculated in **Supplemental Appendix S2**) relative to net growth rate of BaF3s. The distribution is roughly exponential.

Allele-specifc mutation probabiltiies $rho_i$ were assigned from a uniform distribution and normalized to sum to unity. The resistance mutation rate was $\mu = 10^{-8}$ /division.

Tumor size upon detection/resection was variable, taking on values $M_{pre} = [10^9, 10^{10}, 10^{11}, 10^{12}]$ cells. Likewise, the population size after resection took on values $M_{post} = [10^5, 10^6, 10^7, 10^8]$ cells.

#### 0.3 Model Details

Cancer evolution was modeled stochastically as a continuous-time birth-death-mutation process. Simulations were executed in Matlab using a Gillespie algorithm. The propensity vector $a(\mathbf{x})$ and state change matrix $\nu$ were defined as follows:

$$a(\mathbf{x}) = \frac{1}{a_0} \begin{bmatrix} b(1-\mu)S \\ (d+\alpha_S)S \\ b\rho_1\mu S \\ bR_1 \\ (d+\alpha_1)R_1 \\ b\rho_2\mu S \\ bR_2 \\ \vdots \end{bmatrix} \tag{1}$$

$$\nu = \begin{bmatrix} 1 & 0 & 0 & \dots \\ -1 & 0 & 0 & \dots \\ 0 & 1 & 0 & \dots \\ 0 & 1 & 0 & \dots \\ 0 & -1 & 0 & \dots \\ 0 & 0 & 1 & \dots \\ 0 & 0 & 1 & \dots \\ \vdots & \vdots & \vdots & \ddots \end{bmatrix} \tag{2}$$

Here, $a_0$ is the sum of the rates in the propensity function.

Upon detection, the sensitive population was reduced to $M_{post}$. If any resistant cells were present prior to resection, their population size after resection was drawn from a binomial distribution with probability $p = \frac{M_{post}}{M_{pre}}$. Before detection, $\alpha_i = 0$ for all populations. After detection, $\alpha_i$ values were updated with appropriate drug kill terms for each variant.
